# Forward genetic screen in zebrafish identifies new fungal regulators that limit host-protective *Candida*-innate immune interaction

**DOI:** 10.1101/2025.02.14.638315

**Authors:** Bailey A. Blair, Emma Bragdon, Gursimran Dhillon, Nnamdi Baker, Lena Stasiak, Mya Muthig, Pedro Miramon, Michael C. Lorenz, Robert T. Wheeler

**Affiliations:** Department of Molecular & Biomedical Sciences, University of Maine, Orono, ME 04469; Graduate School of Biomedical Sciences and Engineering, University of Maine, Orono, ME 04469; Department of Microbiology and Molecular Genetics, McGovern Medical School, The University of Texas Health Science Center at Houston, Houston, USA

## Abstract

*Candida* is one of the most frequent causes of bloodstream infections, and our first line of defense against these invasive infections is the innate immune system. The early immune response is critical in controlling *C. albicans* infection, but *C. albicans* has several strategies to evade host immune attack. Phagocytosis of *C. albicans* blocks hyphal growth, limiting host damage and virulence, but how *C. albicans* limits early recruitment and phagocytosis in vertebrate infection is poorly understood. To study innate immune evasion by intravital imaging, we utilized the transparent larval zebrafish infection model to screen 131 *C. albicans* mutants for altered virulence and phagocyte response. Infections with each of seven hypovirulent mutants led to altered phagocyte recruitment and/or phagocytosis, falling into four categories. Of particular interest among these is *NMD5*, a predicted β-importin and newly-identified virulence factor. The *nmd5*Δ/Δ mutant fails to limit phagocytosis and its virulence defects are eliminated when phagocyte activity is compromised, suggesting that its role in virulence is limited to immune evasion. These quantitative intravital imaging experiments are the first to document altered *Candida*-phagocyte interactions for several additional mutants, and clearly distinguish recruitment from phagocytic uptake, suggesting that *Candida* modulates both events. This initial large-scale screen of individual *C. albicans* mutants in a vertebrate, coupled with high-resolution imaging of *Candida*-phagocyte interactions, provides a more nuanced view of how diverse mutations can lead to more effective phagocytosis, a key immune process which blocks germination and drives anti-fungal immunity.

**Importance:** *Candida albicans* is part of the human microbial community and is a dangerous opportunistic pathogen, able to prevent its elimination by the host immune system. Although *Candida* avoids immune attack through several strategies, we still understand little about how it regulates when immune phagocytes get recruited to the infection site and when they engulf fungal cells. We tested over 130 selected *Candida* mutants for their ability to cause lethal infection and found several avirulent mutants which provoked altered innate immune responses, resulting in lower overall inflammation and greater host survival. Of particular interest is *NMD5*, which acts to limit fungal phagocytosis and is predicted to regulate the activity of stress-associated transcription factors. Our high-content screening was enabled by modeling *Candida* infection in transparent vertebrate zebrafish larva. Our findings help us understand how *Candida* survives immune attack during commensal and pathogenic growth, and may eventually inform new strategies for controlling disease.

## Introduction

*Candida albicans* is the one of most common bloodstream infections in the U.S. causing approximately 25,000 cases annually (CDC). *C. albicans* can normally be found as a commensal in the gastrointestinal tract, mouth, skin, or vagina in up to 70% of the population (1–3). While *C. albicans* is found in healthy individuals it can also cause infections ranging from superficial mucosal infections such as vulvovaginal candidiasis and oropharyngeal candidiasis, to lethal systemic infections with attributable mortality rates of approximately 25% (4, 5). The host immune response is tasked with protecting individuals from these infections with the innate immune system being of special importance in fighting systemic *Candida* infections. In turn, *C. albicans* employs many mechanisms to subvert the actions of the host immune attack (6–14). While we understand some of how *C. albicans* can evade host immune responses *in vitro*, we still know little about this during vertebrate infection.

The innate immune response is the first line of defense against *C. albicans*, and is critical in controlling and preventing systemic candidiasis (15–19). This is highlighted by the fact that patients with neutropenia are more susceptible to invasive *Candida* infections, and mice with macrophage defects survive experimental systemic infection poorly. Phagocytes get to the infection site by following cytokine and chemokine gradients and presumably identify fungal cells for ingestion using fungal-derived chemoattractants (17, 18, 20). While phagocytes play crucial roles, other innate immune cells such as epithelial cells, microglia, natural killer cells and innate lymphocytes also play important roles (18, 21). Cytokines and chemokines, which bring phagocytes to the infection site, simultaneously activate them and induce their differentiation. Once there, phagocytes must locate fungal cells by soluble cues, recognize the foreign microbial cells based on surface patterns and opsonins, and initiate phagocytosis.

Immune cells such as phagocytes recognize pathogen associated molecular patters (PAMPs) in *C. albicans* cell wall, but *C. albicans* is able to shield them from immune cells behind a layer of mannosylated proteins of the outer cell wall (9). Macrophages and neutrophils are the main effector cells against *C. albicans* and employ many strategies to kill *C. albicans.* These cells are able to phagocytose *C. albicans* yeast as well as short hyphae, produce antimicrobial peptides, reactive oxygen species, and extracellular traps to combat *C. albicans* (8, 13, 14, 22). Not only can *C. albicans* shield its cell wall PAMPs from these cells, but once taken up by a phagocyte *C. albicans* can survive by preventing the fusion of the phagosome with the lysosome, alkanizing the acidic environment of the phagolysosome, producing catalase and superoxide dismutase to counteract ROS, and upregulating DNA repair systems and heat shock proteins to counteract damage caused to DNA and proteins (23, 24). In addition, *C. albicans* has also been seen to escape from host cells such as macrophages by inducing pyroptosis; or also, although rare, vomocytosis (25) (23). These mechanisms were initially described *in vitro*, yet we still do not fully understand which mechanisms play critical roles during infection or which fungal pathways mediate these activities The larval zebrafish provides a unique model that is well-suited to investigate the interactions between *C. albicans* and the vertebrate innate immune response (26–28). The transparency and availability of many transgenic lines permits quantitative imaging of the immune response to *C. albicans* infection in the context of a live host. Furthermore, the small size and fertility of zebrafish enables cost-effective moderate- to high-throughput screening in a vertebrate model. Previous results suggest that the early phagocyte response is critical to survive a *C. albicans* hindbrain ventricle infection (29, 30). Evidence from the larval zebrafish also suggest that *C. albicans* has the ability to limit this response be reducing the recruitment of phagocytes to the infection site (30). This ability to limit phagocyte recruitment was observed for a WT *C. albicans* strain, but not a yeast locked strain, suggesting this response may be regulated with the yeast to hyphal transition.

We sought to identify new *C. albicans* factors playing a role in limiting early phagocyte responses by leveraging the transparent zebrafish infection model. Since virulence is linked to early phagocytic efficiency, we screened 131 engineered *C. albicans* mutants for virulence defects in the larval zebrafish hindbrain infection model. Since there may be links between evasion of phagocyte recruitment and the yeast-to-hyphal transition, we chose a set of mutants that had been characterized in a previous high-throughput pooled screen as having either an infectivity defect only or a morphogenesis defect only (31). Since little is known about soluble chemoattractants secreted by *Candida*, we also included single mutants from groups of genes that code for potential secreted proteins such as secreted aspartyl proteases and lipases.

Mutations that were associated with hypovirulence and could be faithfully complemented were then screened for multiple phagocyte recruitment and phagocytosis phenotypes during early infection. Several genes previously known to alter morphology and/or virulence were found to limit early phagocytosis of *Candida*, a previously unknown function of these genes. Strikingly, the predicted karyopherin *NMD5* lost its virulence defect when the host was immunosuppressed in any of three ways—suggesting that its role in virulence is largely confined to limiting early phagocyte recruitment and phagocytosis. These results expand our understanding of how *Candida* virulence genes mediate pathogenesis through limiting the early innate immune response.

## Results

### Forward genetic screen for altered fungal immune evasion based on loss of virulence

*C. albicans* is known to limit immune recruitment and phagocytosis during infection, although morphological switching can regulate phagocyte recruitment, few molecular details are known about how this occurs (30, 32, 33). The zebrafish hindbrain infection model provides a useful *in vivo* system to intravitally image early fungal and host dynamics, and has identified a close correlation between early phagocyte-mediated fungal containment and overall survival (29, 30, 33). We leveraged these advantages to screen individual *C. albicans* mutants for virulence and phagocytosis defects, with an initial screen for hypovirulence and a secondary screen for altered fungal-phagocyte interaction.

We used a small number of mutants to define infection parameters and enable high-throughput screening; these mutants have normal *in vitro* competitive fitness, were present in our strain collections, and are predicted to have cell wall defects (*mnn15*Δ/Δ, *mnt1*Δ/Δ), known to have filamentous growth defects or altered interaction with phagocytes *in vitro* (*mad2*Δ/Δ, *ece1*Δ/Δ, *pra1*Δ/Δ), and/or hypovirulence in murine models (Table S1) (34–37). In initial virulence tests, two mutant strains were tested along with controls and at least 3 biologically independent experiments were performed with approximately 50 fish infected per mutant (Fig. 1A). Inoculums were counted by fluorescence microscopy to ensure they received the correct amount of *Candida* (10-25 fungal cells), then larvae were followed for survival for three days relative to the SN250 wildtype (Fig. 1B). Three of the nine strains tested had significantly reduced (*ssu81*Δ/Δ & *mad2*Δ/Δ) or abolished virulence *rbt1*Δ/Δ*^968-2166^* (Fig. 1C). We then used the average and standard deviation of 72 hours post infection (hpi) survival for wildtype-infected fish to determine z-score cutoffs for subsequent experiments, to exclude data in which wildtype-infected survival was out of range (average +/- 2.5 SD [20 - 80% survival]). In addition, we quantified host-pathogen interactions by confocal microscopy at 4-6 hpi, scoring fungal cells as intra-versus extra-cellular based on a combination of Calcofluor white staining of the inoculum and DIC imaging of host phagocytes (Fig. 1D-insets & 1E). Although this method was limited because only the initial inoculum was fluorescently stained and phagocytes were not fluorescent, there was a consistent trend for increased fungal phagocytosis of *mad2*Δ/Δ compared to the control SN250 (Fig. 1E, Table S2, p=0.009, effect size = 0.90; large). The *rbt4*Δ/Δ mutant was phagocytosed significantly less efficiently (p=0.029, effect size =0.55; large), but was not pursued further because the lower phagocytosis was not associated with altered survival (Fig. 1C). Interestingly, *mad2Δ/Δ* was also one of three strains with significantly reduced virulence (Fig. 1B-C). The other two hypovirulent mutants failed later validation steps—*rbt1*Δ/Δ*^968-2166^*failed at the complementation step and *ssu81*Δ/Δ failed when the second isolate was tested. Although morphological quantification of the fungi was not possible because only the inoculum was labeled, and the inoculum was all yeast cells for all strains, there were no qualitative differences noted in the amount of filamentous growth (as visualized by brightfield microscopy) between SN250 and any of the mutant strains.

**Fig. 1.**
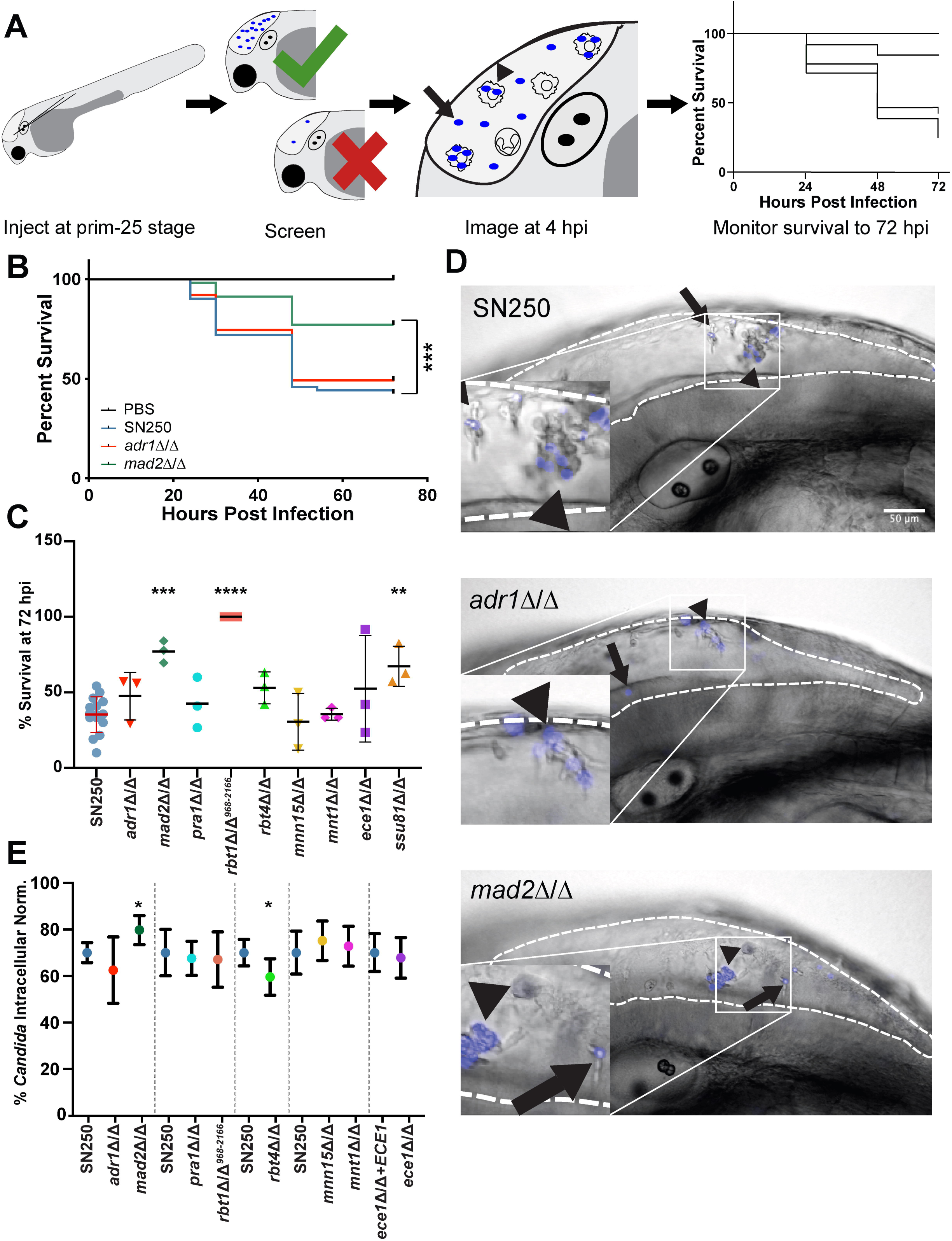
Defining infection parameters. **A)** Flow chart showing workflow of pilot experiments. Hindbrain infections were performed at the prim-25 stage, and fish were then screened to ensure they received the correct inoculum (10-25 cells). At 4-6 hours post-infection, fish were imaged by confocal microscopy to score fungal phagocytosis; survival was monitored out to 72 hpi. **B)** Example Kaplan-Meier survival curves pooled from 3 experiments showing fish injected with PBS (Control, n=83), SN250 (WT, n=61), *adr1*Δ/Δ (n=63), or *mad2*Δ/Δ (n=57). Fish injected with *mad2*Δ/Δ showed increased survival compared to SN250 (p=0.0001). **C)** Survival of fish injected with each strain at 72 hpi in three independent experiments. Individual points represent biologically independent experiments on different days. Bars show means and standard deviations with SN250 in red to depict WT cutoff range for inclusion of experiments. Significant differences in survival curves were determined by Mantel-Cox log rank tests comparing the mutant strain to SN250 from data pooled from three biological replicates of the same experiments. Two mutants were tested per experiment, and Bonferroni corrections were performed. **D)** Representative images of hindbrain ventricle infection to score fungal phagocytosis at 4 hpi. *C. albicans* initial inoculum was stained with Calcofluor white, shown in blue. The hindbrain ventricle is outlined by a white dashed line. Scalebar is 50 μm. Arrows point to extracellular *Candida*, while arrowheads point to intracellular *Candida*. **E)** Quantification of the percent of intracellular *Candida*. Fungal cells were scored as intracellular or extracellular from z-stack slices (using Calcofluor fluorescence for the fungi and differential interference contrast (DIC) for imaging the phagocytes) of individual fish taken at 4-6 hpi for each strain. Based on at least 19 fish from at least 3 independent experiments. Significance and effect size were determined as described in Materials & Methods and based on (84). * p<0.05, ** p<0.01, *** p<0.001.

A total of 131 mutant *C. albicans* strains with expected deficiencies in predicted secreted factors, hyphal growth, or virulence were then selected for screening (Table S1), based on their phenotypes observed in previous screens (31). One group of strains (the Morphology category) were selected for a published defect in hyphal growth on Spider medium but no defect in virulence, as we hypothesized that these mutants might disrupt the co-regulation of immune evasion mechanisms with the yeast-to-hyphal transition (10). While these strains have a morphogenesis defect on Spider plates, defects in filamentous growth are often very dependent on the environmental context and strain, and therefore may or may not have a filamentous growth defect in the zebrafish hindbrain (38–40). A complementary set of strains (the Infectivity category) had a competitive defect in pooled mouse infection but no morphogenesis defect on Spider agar; we reasoned these strains may be cleared more effectively by the host immune response even if they are not defective in filamentous growth. This included 69 mutants that had a morphogenesis defect on Spider agar but no pooled virulence defect, 41 that had an infectivity defect in pooled infection but no Spider morphogenesis defect, and one had both defects. The final category (Secreted-Predicted) included 20 genes encoding predicted secreted peptides— including lipases, proteases and other genes annotated as potentially secreted—but the mutants had no Spider agar morphogenesis or pooled virulence defect (31) (Table S1).

In this primary screen we chose to facilitate high-throughput screening for cell-autonomous virulence defects, so inoculums were not counted and no replicates were performed. Virulence testing revealed several mutants with greatly reduced virulence, as measured by z-score (based on deviance from the mean for WT infection, see Materials and Methods). Seventeen had a fish survival z-score > 3, while 27 had a z-score between 2 and 3 (Fig. 2). Of the 41 strains in the Infectivity category, 6 of these had a z-score >3, with another 6 between z-scores of 2 and 3. Out of the 70 in the Morphogenesis category, 11 had z-score > 3, with another 16 between 2 and 3. In addition, 4 genes from the secreted aspartyl protease (SAP) family of genes had z-scores between 2 and 3. As these fish were not screened to ensure the correct number of *C. albicans* injected (10-25 fungal cells), we first retested hypovirulent strains with z-scores > 3 with an added step of screening for inoculum per fish. On retest, both independent isolates from the Noble library were tested and strains were genotyped to confirm the correct gene deletion. After retesting, this led to a total of 10 mutants with reproducible hypovirulence: *rbt1*Δ/Δ*^968-2166^, orf19.5547*Δ/Δ, *pep8*Δ/Δ*, cht2*Δ/Δ, *apm1*Δ/Δ, *rim101*Δ/Δ, *brg1*Δ/Δ, *nmd5*Δ/Δ, *mad2*Δ/Δ, and *cek1Δ/Δ* (Table S1). Hypovirulent strains were then complemented to assess if complementation restored virulence. When available, *in vitro* phenotypes (e.g. morphogenesis defect on Spider media) were also used to assess functional complementation of strains prior to assessing virulence in hindbrain infection. Complementation successfully restored at least some virulence to *brg1*Δ/Δ, *pep8*Δ/Δ*, nmd5*Δ/Δ, *rim101*Δ/Δ, *cek1Δ/Δ*, *apm1Δ/Δ*, and *mad2Δ/Δ* mutants (Fig. 3). It also partially restored *in vitro* filamentous growth and pH-dependent filamentation phenotypes for *brg1*Δ/Δ, *pep8*Δ/Δ and *rim101*Δ/Δ (Fig. S1). We were not able to generate complemented strains that restored even partial virulence to *cht2*Δ/Δ, *orf19.5547*Δ/Δ, or *rbt1*Δ/Δ*^968-2166^* (Fig. S2A-C). Consistent with the failure to complement the partial ORF deletion in *RBT1*, an independently-created full deletion of *RBT1* in the SN250 background did not cause a virulence defect (Fig. S2D). The failure to complement the virulence defects in these strains with the full-length gene suggests that the virulence defect is due to other, non-targeted, genomic changes sustained during their original construction. Mutants that could be complemented were then transformed with pENO1-iRFP (41) to drive cytosolic expression of a near-infrared fluorescent protein for intravital imaging of infections. At the conclusion of this first part of the screen, we were left with seven mutants whose virulence defects could be at least partially complemented with add-back of a full-length copy of the gene: five in the Morphogenesis class (*brg1*Δ/Δ, *pep8*Δ/Δ*, rim101*Δ/Δ, *apm1Δ/Δ*, and *cek1Δ/Δ*), two in the Infectivity class (*nmd5*Δ/Δ and *mad2Δ/Δ*) and none in the Predicted Secreted class.

**Fig. 2.**
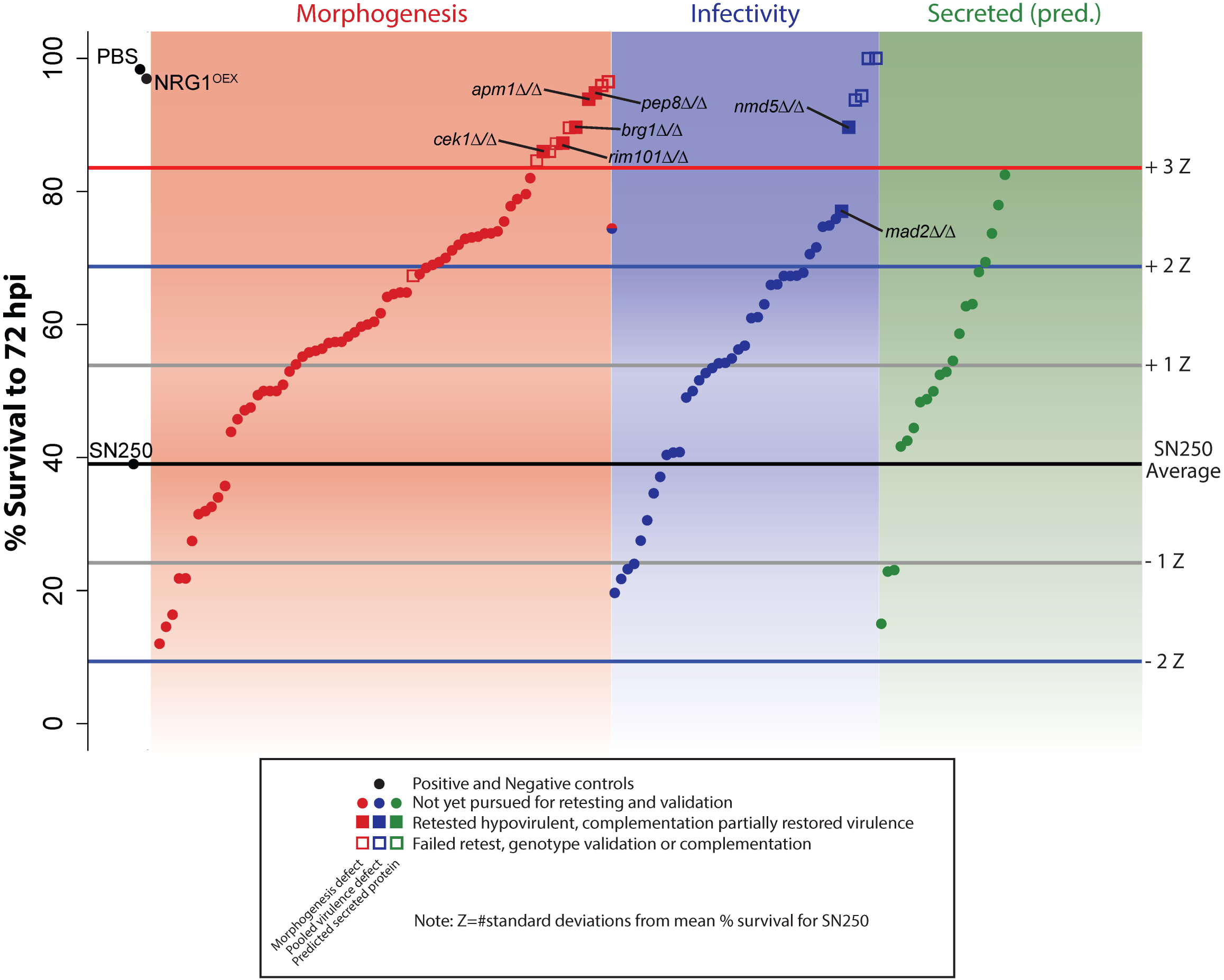
High-throughput virulence screening. Average survival of fish infected with individual mutant *C. albicans* strains (n ≈ 50 fish per mutant strain). Mock infected (PBS) and NRG1^OEX^ infected fish were included as controls. The average survival of the WT SN250 strain is shown by the black line, while differential survival was measured by Z-score (based on the standard deviation of % survival in over 20 experiments with SN250 control infections). Gray lines show Z-score = 1, blue lines show Z-score = 2, and red Z-score = 3. Strains in the red panel were previously seen to have a morphogenesis defect on spider agar, while those in the blue panel showed a defect in pooled virulence tests, and those in the green panel code for predicted secreted proteins. Mutant strains that had a z-score of over 3 were passed to the next phase of screening, shown as squares. Those where both independent mutants showed hypovirulence genotyped correctly, and complementation restored virulence, are shown as filled in squares and were passed to the imaging phase of screening. Those that did not pass secondary screening are shown as empty squares. Complete data is found in Supplementary Table S1.

**Fig. 3.**
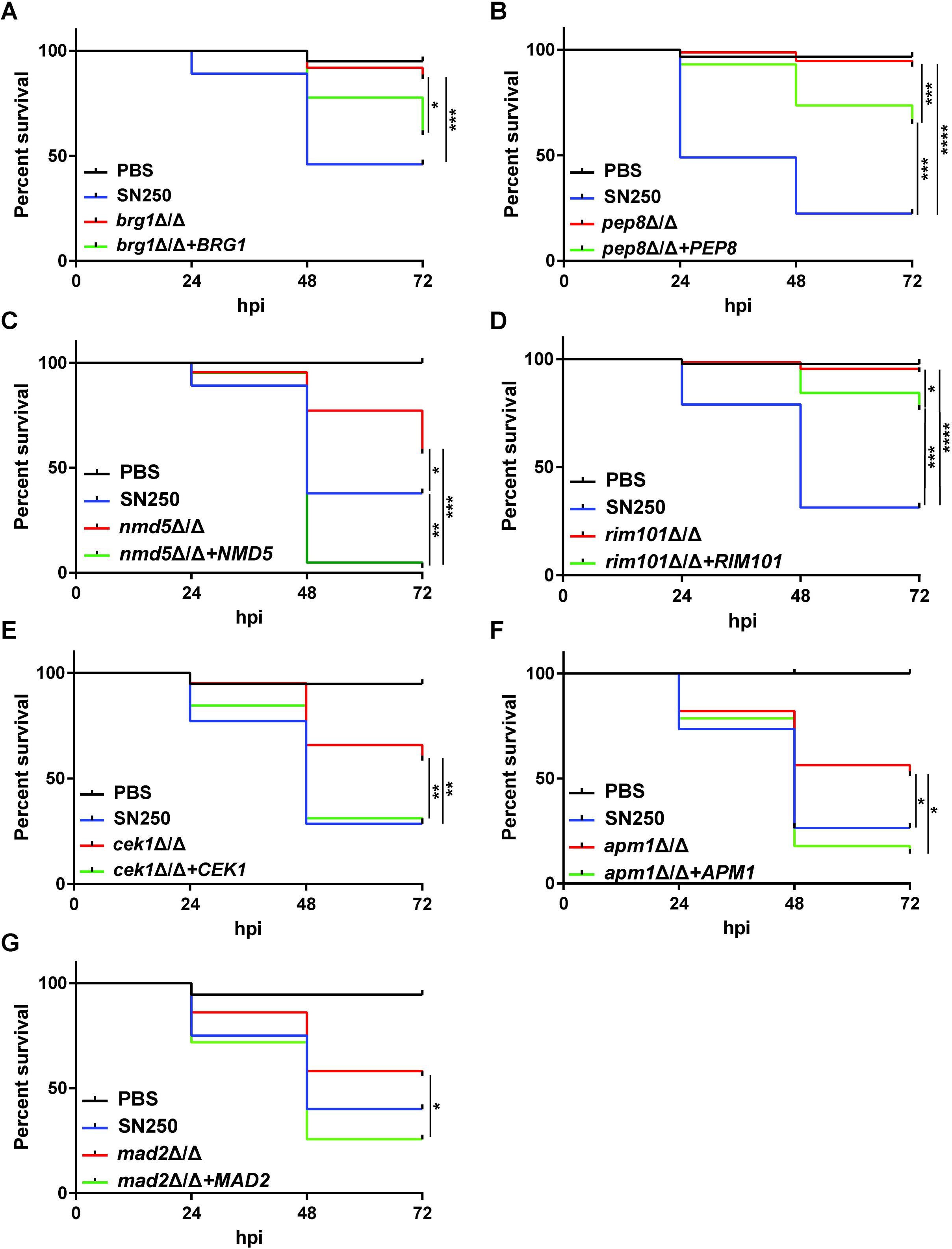
Complementation restores virulence to hypovirulent *C. albicans* mutants. Kaplan-Meier survival curves show restoration of virulence of with complementation. All data in survival curves are pooled from 2 experiments unless otherwise noted. **A)** Fish injected with SN250 (WT, n=37), *brg1*Δ/Δ (n=37), *brg1*Δ/Δ+*BRG1* (n=45), PBS (mock, n= 20). Complementation of *brg1*Δ/Δ restores some virulence. **B)** Fish injected with SN250 (WT, n=49), *pep8*Δ/Δ (n=75), *pep8*Δ/Δ+*PEP8* (n=57), or PBS (mock, n= 30). Complementation of *pep8*Δ/Δ restores some virulence (data pooled from 3 experiments). **C)** Fish injected with PBS (mock, n=20), SN250 (WT, n=37), *nmd5*Δ/Δ (n=44), *nmd5*Δ/Δ+NMD5 (n=41). Complementation significantly increases virulence of *nmd5*Δ/Δ. **D)** Fish injected with SN250 (WT, n= 45), *rim101*Δ/Δ (n=43), *rim101*Δ/Δ+*RIM101* (n=41), or mock infected fish (PBS, n=20). Complementation of *rim101*Δ/Δ restores virulence (Data pooled from 3 independent experiments). **E)** Fish injected with SN250 (WT, n=35), *cek1*Δ/Δ (n=41), *cek1*Δ/Δ+*CEK1* (n=45), or mock infected fish (PBS, n= 19). Complementation of *cek1*Δ/Δ restores virulence. **F)** Fish injected with PBS (mock, n=21), SN250 (WT, n=34), *apm1*Δ/Δ (n=39), or *apm1*Δ/Δ+*APM1* (n=28). Complementation significantly increases virulence of *apm1*Δ/Δ. **G)** Fish injected with PBS (mock, n=18), SN250 (WT, n=40), *mad2*Δ/Δ (n=43), or *mad2*Δ/Δ+*MAD2* (n=39). Complementation significantly increases virulence of *mad2*Δ/Δ. * p_adj_ < 0.05, **, p_adj_ < 0.01, ***, p_adj_ < 0.001

### Altered early phagocyte responses to hypovirulent *Candida* mutants

Previous work has linked efficient early immune phagocytosis of fungi to enhanced survival (29, 30). To determine if the virulence defects for these mutants were associated with a more effective early immune response, we imaged *Tg(mpeg1:GFP)/(lysC:dsRed)* larvae (green macrophages and red neutrophils) infected with iRFP-expressing *Candida* at 4-6 hpi. From these images, we assessed the number of macrophages and neutrophils responding rapidly to infection as well as their ability to phagocytose *Candida*, as measured by the number of extracellular fungi, percent phagocytosis and the number fungi/recruited phagocyte. We chose these measures to quantify (1) the overall ability of phagocytes to internalize fungi, keep them internalized, and prevent their extracellular proliferation; (2) the relative efficiency of phagocytosis, without consideration of the total number of extracellular cells, indicating the overall capacity of recruited phagocytes to engulf fungi; and (3) the average ability of any given phagocyte at the infection site to engulf fungi, providing an indicator of the activation state of the phagocytes and their ability to identify and engulf fungi.

Overall, we found altered acute immune responses to each of the seven validated hypovirulent mutants, with mutant phenotypes in four groups based on infection site immune cell counts and phagocytosis efficiency at 4-6 hpi (Table 1). Three mutants in Group I (*mad2Δ/Δ*, *rim101*Δ/Δ and *brg1*Δ/Δ) were phagocytosed more effectively and there were lower phagocyte numbers at the infection site at 4-6 hpi. Two mutants in Group II (*pep8*Δ/Δ and *apm1Δ/Δ*) had unchanged phagocytosis efficiency and fewer immune cells. One (Group III; *nmd5*Δ/Δ) had greater phagocytosis with an unchanged phagocyte number and one (Group IV; *cek1Δ/Δ*) had increased phagocytosis and an increased phagocyte count. The phenotypic scoring is described below in more detail.

**Table 1:**
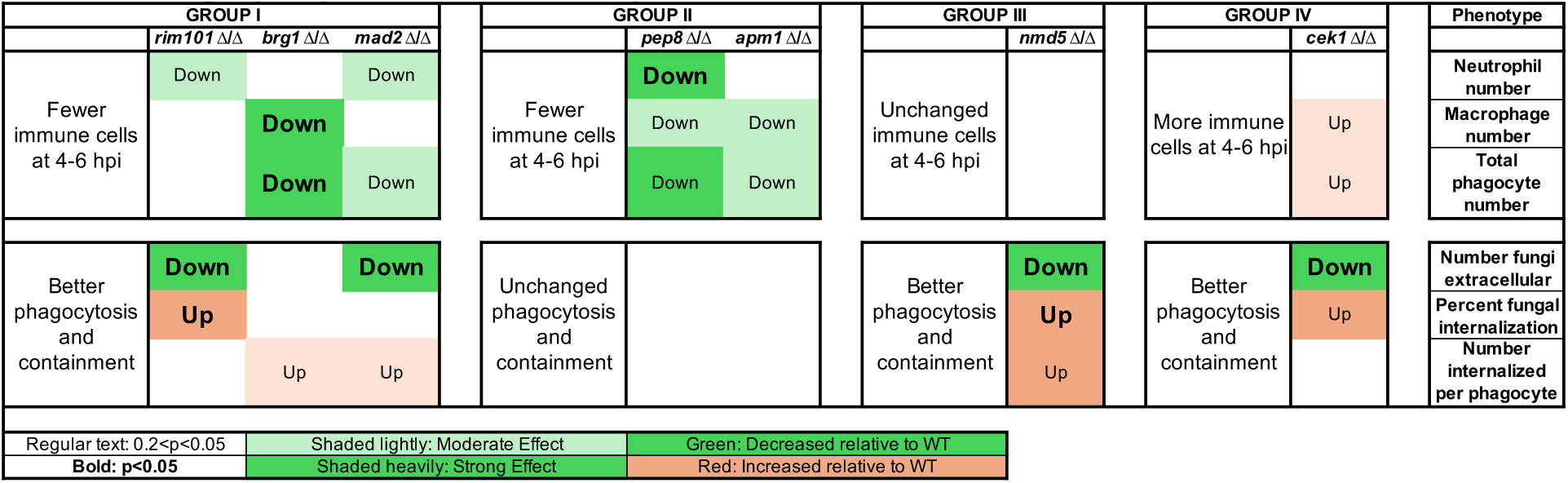
_Mutants grouped by altered innate immune response_.

Mutants in Groups I, II and IV had altered numbers of phagocytes at the infection site at 4-6 hpi, as evidenced by comparisons with wildtype infections (Fig. 4; p-values and effect sizes summarized in Table 1 and detailed in Table S3; see Materials & Methods). Infections with the *cek1*Δ/Δ mutant elicited a higher number of total phagocytes and macrophages (p-values 0.13, 0.14; effect sizes moderate 0.43, 0.41). In contrast, there was a lower number of macrophages at the infection site in *brg1*Δ/Δ and *pep8*Δ/Δ infections (p-values 0.01, 0.10; effect sizes large, 0.69, or moderate, 0.47) and a lower number of neutrophils in *mad2*Δ/Δ, *rim101*Δ/Δ and *pep8*Δ/Δ infections (p-values 0.066, 0.113, 0.046; effect sizes moderate, 0.44, 0.45, or large, 0.59). The small number of neutrophils present early during infection (often zero) suggests that they usually play a limited role early on, so differential neutrophil recruitment likely has more muted biological consequences. Taken together, these data suggest that there is overall decreased phagocyte recruitment to infections with the Group I (*mad2Δ/Δ*, *rim101*Δ/Δ, *brg1*Δ/Δ) and Group II (*pep8*Δ/Δ and *apm1Δ/Δ*) mutants, and overall increased phagocyte recruitment to *cek1Δ/Δ* (Group IV).

**Fig. 4.**
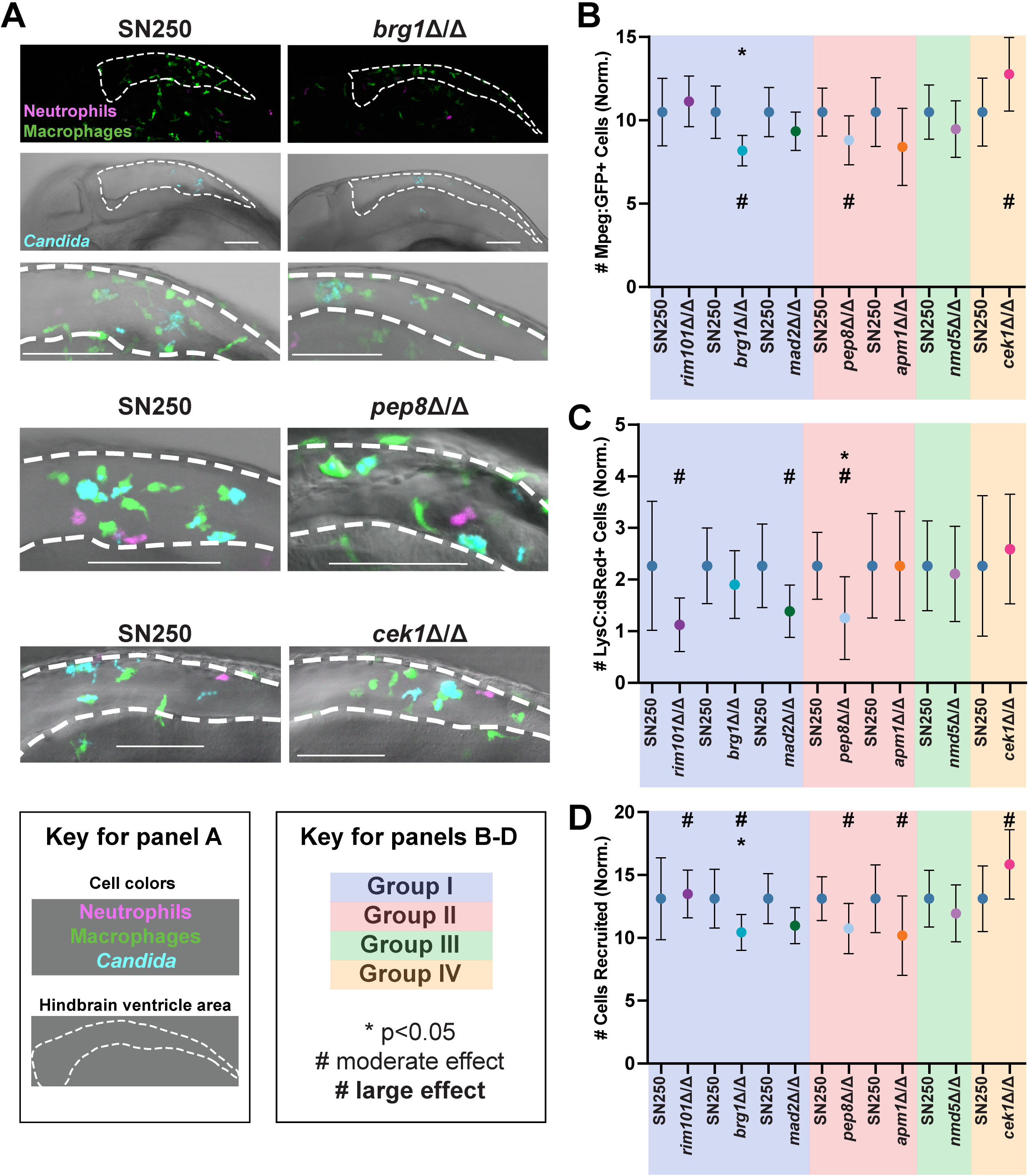
Phagocyte recruitment to hypovirulent *C. albicans* mutants. **A)** Example representative images from *brg1*Δ/Δ- *pep8*Δ/Δ and *cek1*Δ/Δ-infected fish, along with SN250- infected controls, at 4-6 hours post-infection. Images were scored by eye for the number of macrophages (*mpeg1*:GFP+ cells) shown in green and number of neutrophils (*lysC*:dsRed+ cells) in magenta recruited to the infection, as well as if the *Candida* was intracellular or extracellular. Scalebar is 100 μm. **B-D)** Quantification of phagocyte recruitment-related phenotypes. There are separate SN250 columns for each set of experiments, as the mutant was compared to wildtype in the same experiments. **B)** Plots showing the number of *mpeg*:GFP+ macrophages recruited to the infection site normalized to the average amount of *mpeg*:GFP+ macrophages recruited to SN250. **C)** Plots showing the number of *lysC*:dsRed+ neutrophils recruited to the infection site normalized to the average amount of *lysC*:dsRed+ neutrophils recruited to SN250. **D)** Plots showing the number of cells recruited to the infection site normalized to the average recruited to SN250. Cells include *mpeg1*:GFP+ and *lysC*:dsRed+ cells recruited to the hindbrain, as well as non-fluorescent cells containing *Candida*. **(B-D)** Shading indicates the Groups I-IV, based on similar interaction phenotypes (Table 1). Means and 95% confidence intervals are plotted. Statistics were performed from data pooled from at least 3 independent experiments for each mutant, for approximately 25 fish per strain were imaged. Hedges bias-corrected effect sizes and significance was determined for each mutant. * indicates p<0.05, # indicates a moderate effect, while a bold **#** indicates a strong effect.

The ability of phagocytes to engulf fungi—and thereby limit filamentous growth and contain the infection—within the first few hours is associated closely with overall survival of a wildtype infection (29, 30). Mutants in Groups I, III and IV had an overall increase in the ability of phagocytes to internalize fungi. Zebrafish infected with *nmd5*Δ/Δ (Group III) were more effective at each of these measures of fungal internalization (Table 1; Table S3; Fig. 5), with a higher percent internalization (p-value 0.009; large effect size 0.69) and number of fungi per phagocyte (p-value 0.05; large effect size 0.55) and a lower number of extracellular fungi (p-value 0.02; large effect size 0.65). Fish infected with Group I and IV mutants (*mad2Δ/Δ*, *rim101*Δ/Δ, *brg1*Δ/Δ, and *cek1Δ/Δ*) also exhibited at least one measure of increased fungal phagocytosis with at least a moderate effect size (Table 1; Table S3; Fig. 5). Overall, the phagocyte response was able to internalize each of the hypovirulent mutants at least as well as the wildtype strain in the first 4-6 hpi, with five of the seven mutants phagocytosed more effectively than wildtype. This is consistent with the original premise of the screen, which was designed to identify mutants with reduced capacity to avoid innate phagocyte attack by screening initially for hypovirulence.

**Fig. 5.**
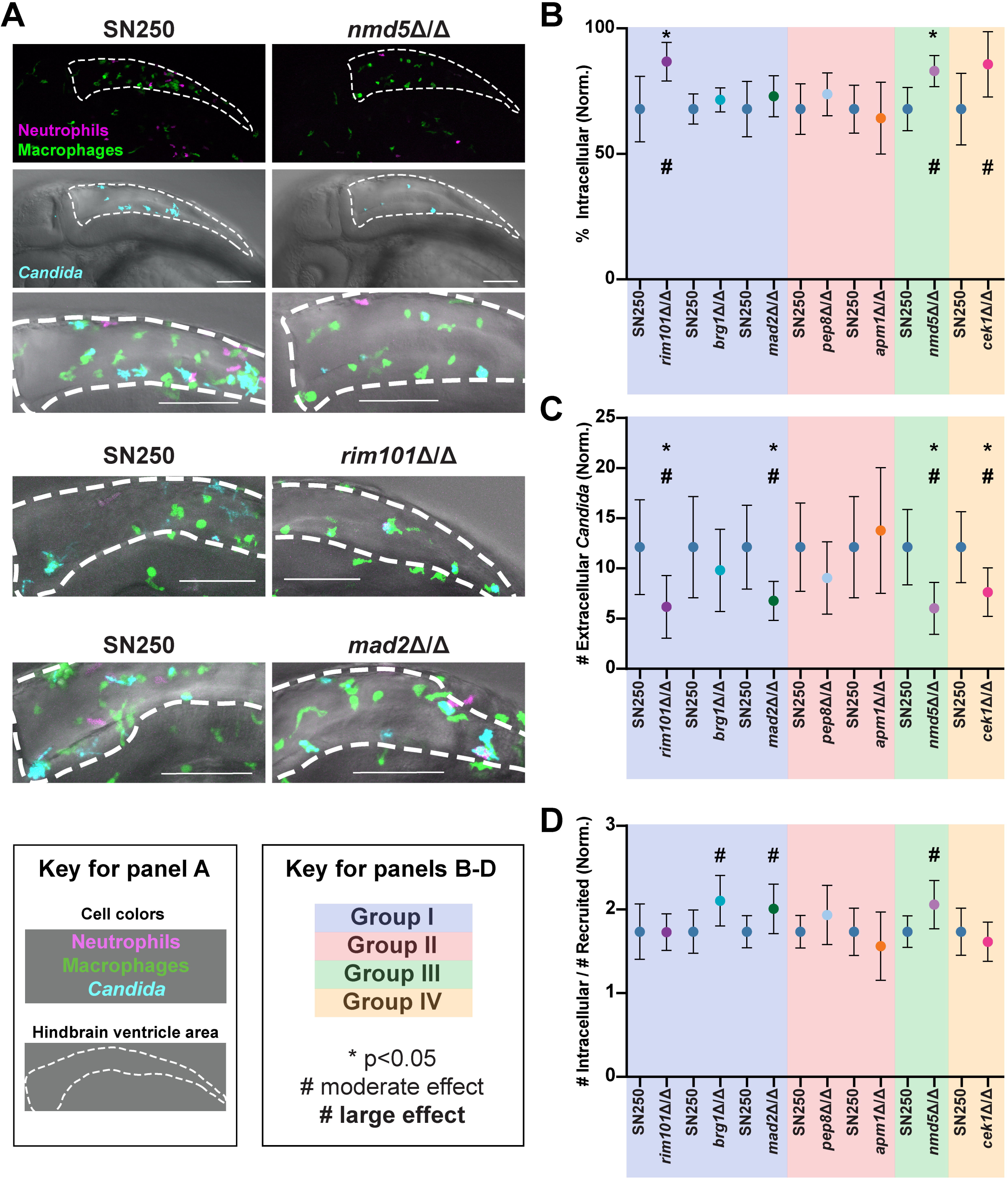
Phagocytosis of hypovirulent *C. albicans* mutants. **A)** Example representative images from *nmd5Δ/Δ- <u>rim101</u>*<u>Δ/Δ</u> and *mad2*Δ/Δ-infected fish, along with SN250-infected controls, at 4-6 hours post-infection. Images were scored by eye for the number of macrophages (Mpeg1-GFP+ cells) shown in green and number of neutrophils (LysC-dsRed+ cells) in magenta recruited to the infection, as well as if the *Candida* was intracellular or extracellular. Scalebar is 100 μm. **B-D)** Quantification of phagocytosis-related phenotypes. There are separate SN250 columns for each set of experiments, as the mutant was compared to wildtype in the same experiments. **B)** Plots of the percent intracellular *Candida* normalized to the average percent intracellular *Candida* for SN250. **C)** Plots showing the number extracellular *Candida* normalized to the average amount for SN250. **D)** Plots showing the number of intracellular *Candida*, divided by the number of cells recruited, normalized to the average for SN250. **(B-D)** Shading indicates the Groups I-IV, based on similar interaction phenotypes (Table 1). Means and 95% confidence intervals are plotted. Statistics were performed from pooled data from at least 3 independent experiments for each mutant, for approximately 25 fish per strain imaged. Hedges bias-corrected effect sizes and significance was determined for each mutant. * indicates p<0.05, # indicates a moderate effect, while a bold **#** indicates a strong effect.

### Fungal morphology defects of mutants early during infection does not correlate with altered innate immune responses

Since filaments are more difficult to phagocytose that yeast, and five of the seven hypovirulent mutants had been previously identified as having filamentous growth phenotypes on Spider agar, we sought to determine if they also had problems switching to filamentous morphology *in vivo* in the first few hours of infection (31, 42–45). We imaged hindbrain infections with each mutant and analyzed the amount of yeast-shaped vs. elongated cells at 4-6 hours post infection, as cells switching to filamentous growth would have had time to grow longer but would not yet have a hyphal shape (Fig. S3A). Not unexpectedly, four of the five mutants with reduced *in vitro* filamentation (*rim101*Δ/Δ, *brg1*Δ/Δ, *pep8*Δ/Δ and *apm1*Δ/Δ, but not *cek1*Δ/Δ) had a reduced number of elongated cells *in vivo* at this early timepoint, with dramatic defects in the *rim101*Δ/Δ, *brg1*Δ/Δ and *pep8*Δ/Δ mutants (Fig. S3B; Table 2, n.b. effect size in table indicated only for those comparisons with p<0.05). On the other hand, neither *cek1*Δ/Δ nor the two mutants in the Infectivity class of mutants (*nmd5*Δ/Δ and *mad2*Δ/Δ) had a significant reduction in filamentous growth. Early phagocytosis is associated with inhibition of germination and lower virulence (29, 30), which could be a factor in reduced filamentous growth. However, since *rim101*Δ/Δ was the only one of the four mutants with reduced filamentous growth that was phagocytosed at a higher rate, this suggests that the other three mutants form fewer filaments *in vivo* because they have an intrinsically reduced ability to switch to filamentous growth during infection. Interestingly, there was no concordance between significantly altered innate immune responses (in recruitment or phagocytosis efficiency) and a reduced ability to switch to filamentous growth in the early hours of infection. For instance, *mad2*Δ/Δ and *rim101*Δ/Δ have very similar phagocyte response profiles (Table 1, Group I), but only *rim101*Δ/Δ has a strong and significant morphogenesis defect at this early time point *in vivo* (Table 2).

**Table 2:** Fungal morphology at 4-6 hpi.

| Table 2: Fungal morphology at 4-6 hpi. |  |  |  |  |  |  |  |
| --- | --- | --- | --- | --- | --- | --- | --- |
|  | GROUP I |  |  | GROUP II |  | GROUP III | GROUP IV |
| Mutant | *rim101 Δ/Δ* | *brg1 Δ/Δ* | *mad2 Δ/Δ* | *pep8 Δ/Δ* | *apm1 Δ/Δ* | *nmd5 Δ/Δ* | *cek1 Δ/Δ* |
| Morphogenesis *in vivo* | Down | Down | NO CHANGE | Down | Down | NO CHANGE | NO CHANGE |
| Regular text: $0.2<p<0.05$ | | Shaded lightly: Moderate Effect | | | Green: Decreased relative to WT | | |
| **Bold: $p<0.05$** |  | Shaded heavily: Strong Effect |  |  | Red: Increased relative to WT |  |  |

**Table 3.** Candida albicans strains.

| Strain | Parental Strain | Genotype | Reference |
| --- | --- | --- | --- |
| *yfgΔ/Δ* (Your Favorite Gene)<br>See Table S1 for complete list of mutant strains | SN152 | *ura3Δ-iro1Δ::imm<sup>434</sup>/URA3-IRO1, his1Δ/his1Δ, arg4Δ/arg4Δ, leu2Δ/leu2Δ, yfgΔ::C.mLEU2/yfgΔ::C.dHIS1 YFG::C.d.ARG4* | (31) |
| SN250-iRFP | SN152 | *ura3Δ-iro1Δ::imm<sup>434</sup>/URA3-IRO1, his1Δ/his1Δ, arg4Δ/arg4Δ, leu2Δ::C.m.LEU2/leu2Δ::C.d.HIS1, , pENO1-iRFP-NATR* | (31), This Study |
| *rbt1Δ/Δ<sup>968-2166</sup>*- iRFP | SN152 | *ura3Δ-iro1Δ::imm<sup>434</sup>/URA3-IRO1, his1Δ/his1Δ, arg4Δ/arg4Δ, leu2Δ/leu2Δ, rbt1Δ<sup>967-2166</sup>::C.mLEU2/rbt1Δ<sup>967-3166</sup>::C.dHIS1, pENO1-iRFP-NATR* | (31), This Study |
| *cht2Δ/Δ*- iRFP | SN152 | *ura3Δ-iro1Δ::imm<sup>434</sup>/URA3-IRO1, his1Δ/his1Δ, arg4Δ/arg4Δ, leu2Δ/leu2Δ, cht2::C.mLEU2/cht2::C.dHIS1, pENO1-iRFP-NATR* | (31), This Study |
| *rim101Δ/Δ*- iRFP | SN152 | *ura3Δ-iro1Δ::imm<sup>434</sup>/URA3-IRO1, his1Δ/his1Δ, arg4Δ/arg4Δ, leu2Δ/leu2Δ, rim101::C.mLEU2/rim101::C.dHIS1, pENO1-iRFP-NATR* | (31), This Study |
| *brg1Δ/Δ*- iRFP | SN152 | *ura3Δ-iro1Δ::imm<sup>434</sup>/URA3-IRO1, his1Δ/his1Δ, arg4Δ/arg4Δ,* *leu2Δ/leu2Δ ,*<br>*brg1::C.mLEU2/brg1::C.dHIS1,*<br>*pENO1-iRFP-NATR* | (31), This Study |
| *cek1Δ/Δ- iRFP* | SN152 | *ura3Δ-iro1Δ::imm<sup>434</sup>/URA3-IRO1,*<br>*his1Δ/his1Δ, arg4Δ/arg4Δ.,*<br>*leu2Δ/leu2Δ ,*<br>*cek1::C.mLEU2/cek1::C.dHIS1,*<br>*pENO1-iRFP-NATR* | (31), This Study |
| *pep8Δ/Δ- iRFP* | SN152 | *ura3Δ-iro1Δ::imm<sup>434</sup>/URA3-IRO1,*<br>*his1Δ/his1Δ, arg4Δ/arg4Δ.,*<br>*leu2Δ/leu2Δ ,*<br>*pep8::C.mLEU2/pep8::C.dHIS1,*<br>*pENO1-iRFP-NATR* | (31), This Study |
| *nmd5Δ/Δ- iRFP* | SN152 | *ura3Δ-iro1Δ::imm<sup>434</sup>/URA3-IRO1,*<br>*his1Δ/his1Δ, arg4Δ/arg4Δ.,*<br>*leu2Δ/leu2Δ ,*<br>*nmd5::C.mLEU2/nmd5::C.dHIS1,*<br>*pENO1-iRFP-NATR* | (31), This Study |
| *apm1Δ/Δ- iRFP* | SN152 | *ura3Δ-iro1Δ::imm<sup>434</sup>/URA3-IRO1,*<br>*his1Δ/his1Δ, arg4Δ/arg4Δ.,*<br>*leu2Δ/leu2Δ ,*<br>*apm1::C.mLEU2/apm1::C.dHIS1,*<br>*pENO1-iRFP-NATR* | (31), This Study |
| *mad2Δ/Δ- iRFP* | SN152 | *ura3Δ-iro1Δ::imm<sup>434</sup>/URA3-IRO1,*<br>*his1Δ/his1Δ, arg4Δ/arg4Δ.,*<br>*leu2Δ/leu2Δ ,*<br>*mad2::C.mLEU2/mad2::C.dHIS1,*<br>*pENO1-iRFP-NATR* | (31), This Study |
| *ece1Δ/Δ - dtom* | BWP17 | *ura3::imm434/ura3::imm434,*<br>*iro1::imm434/iron1::imm434,*<br>*his1::hisG/his1::hisG,*<br>*arg4::hisG/arg4::hisG,*<br>*ece1::HIS2/ece1::ARG4,*<br>*RPS1/rps1::URA3, ENO1/eno1::dTom-*<br>*NATR* | (74) |
| *ece1Δ/Δ+ECE1 - dtom* | BWP17 | *ura3::imm434/ura3::imm434,*<br>*iro1::imm434/iron1::imm434,*<br>*his1::hisG/his1::hisG,*<br>*arg4::hisG/arg4::hisG,*<br>*ece1::HIS2/ece1::ARG4,*<br>*RPS1/rps1::URA3-ECE1,*<br>*ENO1/eno1::dTom-NATR* | (74) |
| *NRG1<sup>OEX</sup>-iRFP* | THE21 | *ade2::hisG::ade2::hisG*<br>*ura3::imm434/ura3::imm434::URA2-*<br>*tetO ENO1/eno1::ENO1 tetR –* | (75, 76) |
|  |  | *ScHAP4AD-3XHA-ADE2 pENO1-iRFP-NATR* |  |
| *rbt1Δ/Δ<sup>968-2166</sup> +RBT1* | *rbt1Δ/Δ<sup>968-2166</sup>* | *ura3Δ-iro1Δ::imm<sup>434</sup>/URA3-IRO1, his1Δ/his1Δ, arg4Δ/arg4Δ., leu2Δ/leu2Δ, rbt1Δ<sup>967-2166</sup>::C.mLEU2/rbt1Δ<sup>967-2166</sup>::C.dHIS1 RBT1::C.d.ARG4* | This Study |
| *rim101Δ/Δ+RIM101* | *rim101Δ/Δ* | *ura3Δ-iro1Δ::imm<sup>434</sup>/URA3-IRO1, his1Δ/his1Δ, arg4Δ/arg4Δ., leu2Δ/leu2Δ, rim101Δ::C.mLEU2/rim101Δ::C.dHIS1 RIM101::C.d.ARG4* | This Study |
| *brg1Δ/Δ+BRG1* | *brg1Δ/Δ* | *ura3Δ-iro1Δ::imm<sup>434</sup>/URA3-IRO1, his1Δ/his1Δ, arg4Δ/arg4Δ., leu2Δ/leu2Δ, brg1Δ::C.mLEU2/brg1Δ::C.dHIS1 BRG1::C.d.ARG4* | This Study |
| *cek1Δ/Δ+CEK1* | *cek1Δ/Δ* | *ura3Δ-iro1Δ::imm<sup>434</sup>/URA3-IRO1, his1Δ/his1Δ, arg4Δ/arg4Δ., leu2Δ/leu2Δ, cek1Δ::C.mLEU2/cek1Δ::C.dHIS1 CEK1::C.d.ARG4* | This Study |
| *pep8Δ/Δ+PEP8* | *pep8Δ/Δ* | *ura3Δ-iro1Δ::imm<sup>434</sup>/URA3-IRO1, his1Δ/his1Δ, arg4Δ/arg4Δ., leu2Δ/leu2Δ, pep8Δ::C.mLEU2/pep8Δ::C.dHIS1 PEP8::C.d.ARG4* | This Study |
| *nmd5Δ/Δ+NMD5-mNeon-iRFP* | *nmd5Δ/Δ-iRFP* | *ura3Δ-iro1Δ::imm<sup>434</sup>/URA3-IRO1, his1Δ/his1Δ, arg4Δ/arg4Δ., leu2Δ/leu2Δ, ndm5Δ::C.mLEU2/nmd5::C.dHIS1 NMD5-mNeon::C.d.ARG4, pENO1-iRFP-NATR* | This Study |
| *apm1Δ/Δ+APM1* | *apm1Δ/Δ* | *ura3Δ-iro1Δ::imm<sup>434</sup>/URA3-IRO1, his1Δ/his1Δ, arg4Δ/arg4Δ., leu2Δ/leu2Δ, apm21Δ::C.mLEU2/apm1Δ::C.dHIS1 APM1::C.d.ARG4* | This Study |
| *mad2Δ/Δ+MAD2* | *mad2Δ/Δ* | *ura3Δ-iro1Δ::imm<sup>434</sup>/URA3-IRO1, his1Δ/his1Δ, arg4Δ/arg4Δ., leu2Δ/leu2Δ, mad2Δ::C.mLEU2/mad2Δ::C.dHIS1 MAD2::C.d.ARG4* | This Study |
| *rbt1Δ/Δ* | SN250 | *ura3Δ-iro1Δ::imm<sup>434</sup>/URA3-IRO1, his1Δ/his1Δ, arg4Δ/arg4Δ., leu2Δ::C.m.LEU2/leu2Δ::C.d.HIS1, rbt1Δ<sup>1-2166</sup>/Δ<sup>1-2166</sup>* | This Study |

**Table 4:** Zebrafish lines.

| Zebrafish Line | Allele | Source/Reference |
| --- | --- | --- |
| AB (Wild Type) | n/a | Zebrafish International Resource Center |
| *Tg(mpeg1:EGFP)/*<br>*Tg(lysC:dsRed)* | gl22Tg<br>nz50Tg | (78, 79) |

**Table 5:**
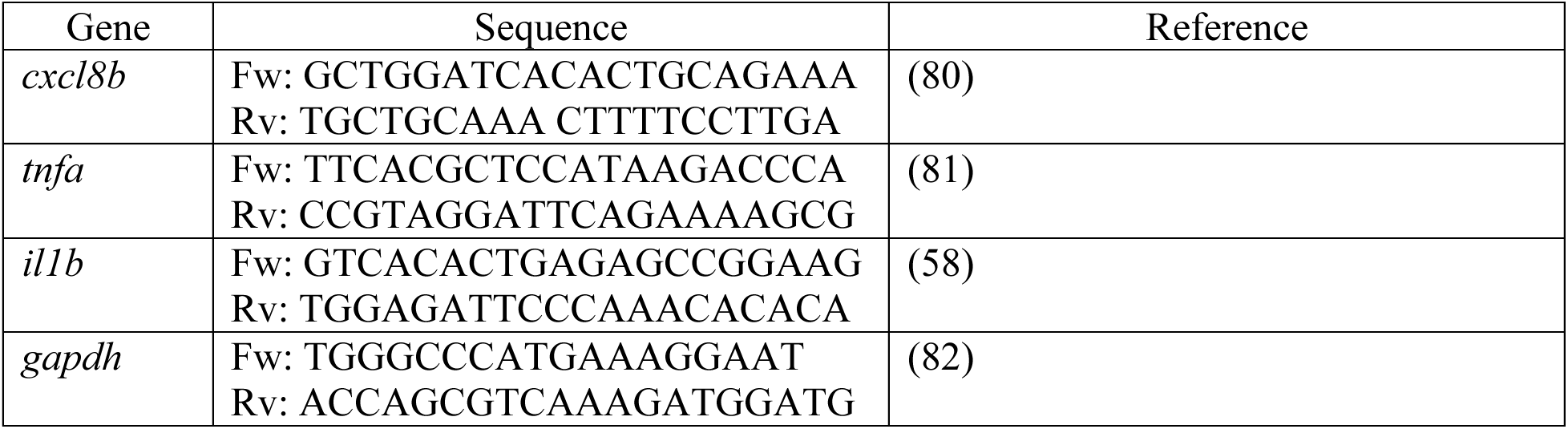
qPCR primers.

### Altered cytokine responses to hypovirulent *Candida* mutants

We reasoned that these altered immune responses might be accompanied by altered expression of proinflammatory cytokines and chemokines. Because immune recruitment and phagocytosis were altered most profoundly for the *brg1*Δ/Δ, *pep8*Δ/Δ and *nmd5*Δ/Δ mutants, and they belong to different classes of mutants (Table 1), we chose to measure inflammatory gene induction in these infections. We measured expression of two key proinflammatory cytokines (interleukin-1 beta and tumor necrosis factor alpha) and the zebrafish IL-8 homolog, each of which is associated with response to *Candida* infection (46). At 4 hours post infection, there was not a significant induction of pro-inflammatory gene expression (Fig. S4), but there was robust induction of these genes by 24 hpi (Fig. 6). At 24 hpi, fish infected with *brg1*Δ/Δ or *pep8*Δ/Δ showed a significant reduction in *cxcl8b*, *tnfa*, and *il1b* induction, while fish infected with *nmd5*Δ/Δ showed a significant reduction in *cxcl8b* and *il1b* induction, but not *tnfa* (Fig. 6A-C). Fish infected with *nmd5*Δ/Δ+*NMD5* showed a trend for increased proinflammatory chemokine/cytokine production, even compared to SN250, which reached significance for *il1b*. This matches well with the decreased survival of fish infected with this complemented strain and the complete complementation phenotype (Fig. 3C). On the other hand, *cxcl8b*, *tnfa*, and *il1b* expression for *brg1*Δ/Δ+*BRG1* and *pep8*Δ/Δ+*PEP8* tended to be between SN250 and the mutant strain, which matches the partial complementation of virulence exhibited by these strains (Fig. 3 A & B). These decreased proinflammatory gene expression signatures are consistent with the effective phagocytosis of the mutants and their overall reduced virulence.

**Fig. 6.**
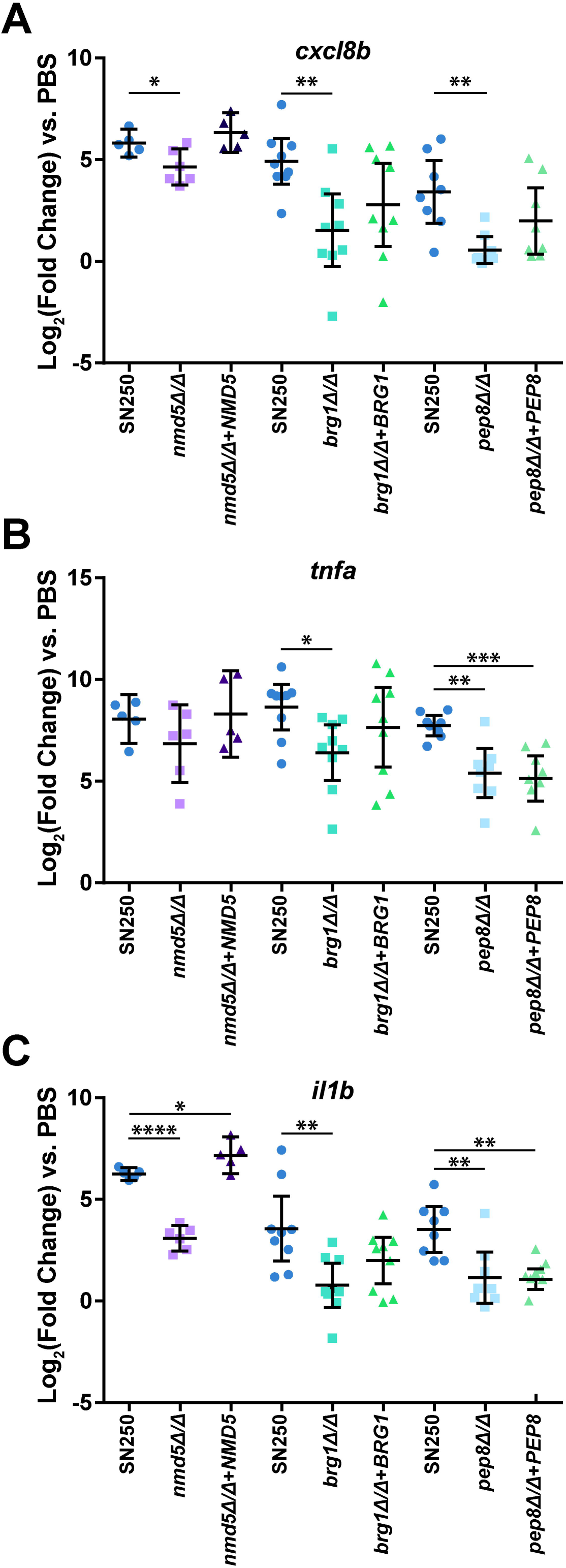
Hypovirulent *C. albicans* mutants elicit a reduced proinflammatory expression at 24 hours post infection. Expression of *cxcl8b* (**A**), *tnfa* (**B**), or *il1b* (**C)** by qPCR analysis of fish infected with WT (SN250), mutant (*nmd5*Δ/Δ, *brg1*Δ/Δ, or *pep8*Δ/Δ), or complemented (*nmd5*Δ/Δ+NMD5, *brg1*Δ/Δ+BRG1, or *pep8*Δ/Δ+PEP8) *C. albicans* at 24 hpi. Each point represents a pool of at least 5 larvae, and data was pooled from 3 (*NMD5*) or 4 (*BRG1* & *PEP8*) independent experiments. Gene expression was normalized to *gapdh* and induction was determined relative to PBS mock infected larvae. Significance was determined by one-way ANOVA with Dunnett’s multiple comparisons tests.

### *NMD5* is only required for virulence in the presence of fully-active immune attack

Given that *nmd5*Δ/Δ infections are associated with greater phagocytosis and thus infection containment, and its role in *C. albicans* has not been previously described, we sought to determine if its primary role in virulence is in immune evasion. In *S. cerevisiae, Sc*Nmd5p is required for transport of *Sc*Hog1p and *Sc*Crz1p into the nucleus and *ScNMD5* mutants are sensitive to salt stress imposed by NaCl, LiCl, MnCl_2_, and CaCl_2_ (47–49). We therefore tested if *C. albicans nmd5*Δ/Δ was also sensitive to salt stress, oxidative stress or pH. While *nmd5*Δ/Δ formed smaller colonies on regular YPD plates, its relative ability to grow on YPD was unchanged with any of these stresses (Fig. S5).

We reasoned that if the virulence defect that we observed for *nmd5*Δ/Δ was due to a failure to evade phagocytosis, then limiting the immune response should enhance virulence of this mutant. We tested this in several ways. First, we treated infected fish with dexamethasone, a general immunosuppressant that regulates macrophage activity in zebrafish (50). As expected based on previous results, *nmd5*Δ/Δ was less virulent than SN250 in the control DMSO/vehicle treatment condition (Fig. 7A, p=0.014). Dexamethasone immunosuppression increased the virulence of *nmd5*Δ/Δ and eliminated the difference in virulence between *nmd5*Δ/Δ and its wildtype SN250 control (Fig. 7A, p_adj_<0.0006). We then selectively inactivated NADPH oxidase, knocking down p47*^phox^*, to reduce phagocyte recruitment to and phagocytosis of *C. albicans* (30). As expected based on previous results, *nmd5*Δ/Δ was less virulent than SN250 in the control STD morpholino condition and SN250 was more virulent in the p47*^phox^* morphant fish as compared to the STD control morphants (Fig. 7B; p=0.0002 and p=0.039, respectively). This gene-directed inactivation caused *nmd5*Δ/Δ to become more virulent (p_adj_=0.0012), and eliminated any difference in survival between SN250- and *nmd5*Δ/Δ-infected fish (Fig. 7B). Lastly, we performed yolk infections, as there is a weaker immune response to yolk infection relative to hindbrain infection (51). In yolk infections, we also observed no significant difference in the virulence of *nmd5*Δ/Δ compared to SN250 (Fig. 7C). These three models of reduced immune response/immunosuppresssion consistently show that *nmd5*Δ/Δ is just as virulent as wildtype SN250 when innate immunity is limited, suggesting that its lack of virulence is due to its failure to evade the immune response.

**Figure 7.**
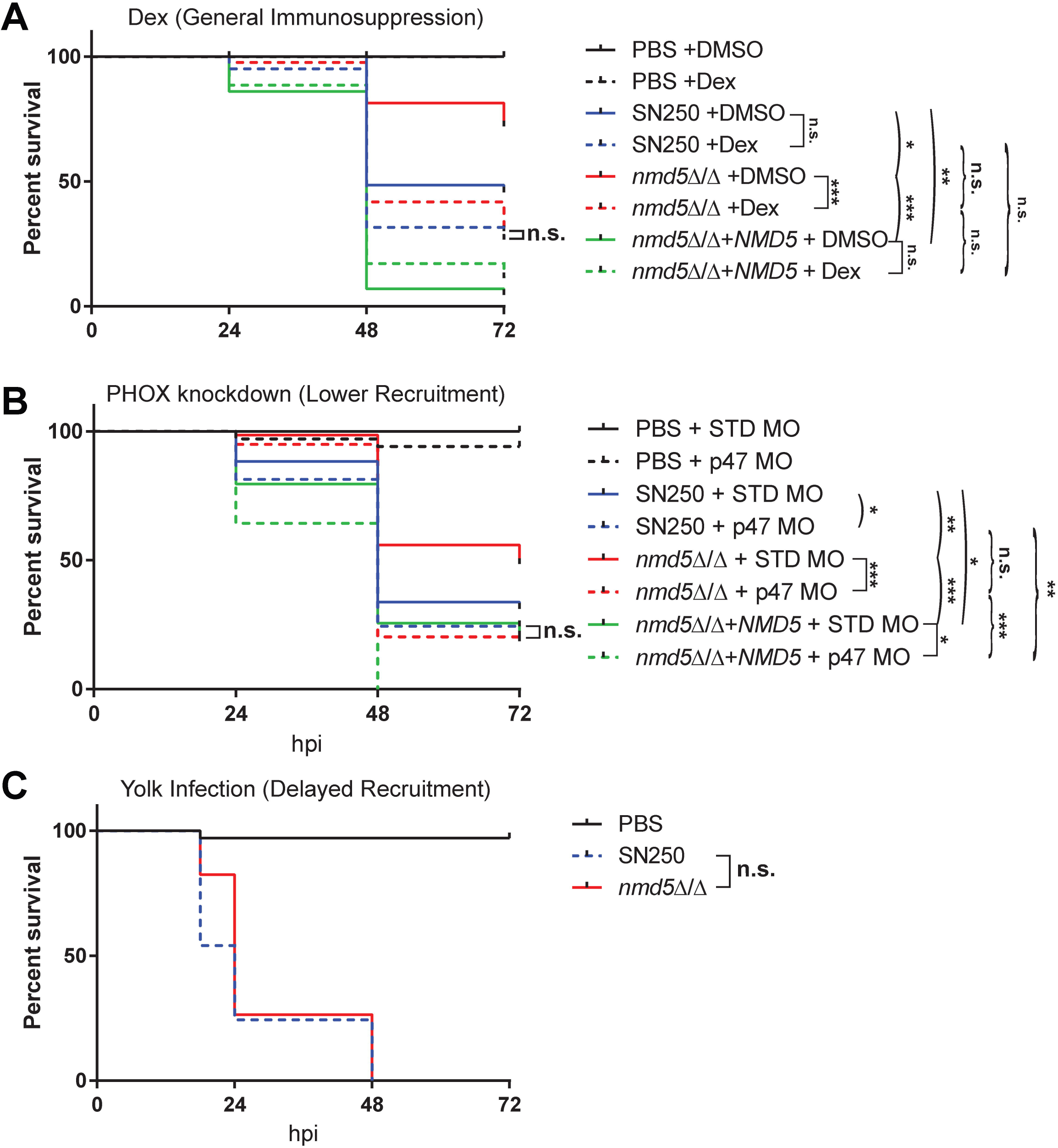
n*m*d5Δ/Δ has fully restored virulence in fish with a reduced immune response. **A)** Kaplan-Meier survival curve of dexamethasone treated hindbrain injected fish with PBS (n=28), SN250 (n=41), *nmd5*Δ/Δ (n=43), or *nmd5*Δ/Δ+*NMD5* (n=35) or DMSO fish injected with PBS (n=28), SN250 (n=36), *nmd5*Δ/Δ (n=43), or *nmd5*Δ/Δ+*NMD5* (n=43). Data pooled from 3 independent experiments. **B)** Kaplan-Meier survival curve of standard morphant fish injected with PBS (n=22), SN250 (n=46), *nmd5*Δ/Δ (n=39), or *nmd5*Δ/Δ+*NMD5* (n=35), and p47 morphant fish injected with PBS (n=20), SN250 (n=59), *nmd5*Δ/Δ (n=26), or *nmd5*Δ/Δ+*NMD5* (n=34). Data pooled from 4 independent experiments. **C)** Kaplan-Meier survival curve of PBS (n=34), SN250 (n=37), or *nmd5*Δ/Δ (n=34) yolk injected fish. Data pooled from 2 independent experiments. Statistics were performed as described in detail in Materials & Methods. Pairwise comparisons that are shown by arcs are confirmatory based on previous experiments; those shown by brackets are exploratory and adjusted for multiple comparisons by Bonferroni correction. Square brackets show if there is an effect of an immune perturbation on survival; curved brackets show if there is an effect of genotype on survival in the context of an immune perturbation. N.s. p_adj_ >0.05 * p_adj_ < 0.05, **, p_adj_ < 0.01, ***, p_adj_ < 0.001.

## Discussion

*Candida albicans* has evolved over many generations with vertebrate hosts and has developed the ability to avoid immune clearance through activities such as filamentous growth, masking of cell wall epitopes, production of a toxin and avoidance of antibody opsonization (6–14). However, we still know little about how each of these abilities affects immune evasion during vertebrate infection and we know even less about which fungal genes and pathways regulate immune evasion. The transparency of the larval zebrafish model is a powerful tool that can be utilized to elucidate the different mechanisms of immune evasion in *C. albicans*, especially combined with its cost-effectiveness. Previous work in this infection model has shown that differential immune recruitment and fungal containment through phagocytosis represent important predictors for the fate of individual hosts (29, 30). These favorable aspects of the model led us to complete the first medium-scale screen of 131 *C. albicans* mutants for virulence, with subsequent analysis for early immune-mediated fungal phagocytosis. This screen characterized several new and known virulence genes as having previously unknown roles in limiting innate immune responses at the infection site. We also identified *NMD5* as a new virulence factor that enables immune evasion.

This is the first single-mutant infection screen of more than 100 individual *C. albicans* mutants in any vertebrate infection model, made possible using a zebrafish model. Very few virulence screens of more than 100 *C. albicans* mutants have been conducted, all using pooled/barcode screening methodology (31, 52, 53). While this is a powerful method, secreted signals and virulence factors that affect the overall environment of the infection site will be missed due to the majority prevalence of cells lacking the phenotype. Pooled virulence screens also score competitive index rather than virulence, per se, potentially missing virulence factors. We chose a zebrafish larval hindbrain infection model to screen individual mutants because it overcomes these drawbacks, reproduces many aspects of murine disseminated infection, and provides a useful infection route for quantifying phagocyte recruitment and response (27, 30, 54). As expected, due to the limitations of pooled screening, several mutants that are hypovirulent in both murine tail vein infection and in our screen were missed in the previous pooled screens (*rim101*Δ/Δ, *brg1*Δ/Δ, *cek1*Δ/Δ).

Of the seven mutants identified here, only four mutants—in *RIM101*, *BRG1*, *MAD2*, *CEK1*—have been tested in single-strain murine tail vein infections; all of them are hypovirulent (Table S4) (44, 55–57). Pooled screens have identified mild defects in competitive index for each of these other three mutants (31, 42). All of the mutants with intermediate virulence defects in our model that have been tested in mouse tail vein infection are also hypovirulent, suggesting that this class represents a mine of new virulence genes, many of which are uncharacterized with only an ORF number (Table S4). However, the converse is not true and some mutants hypovirulent in the mouse were not hypovirulent in our study, suggesting that there may be some murine-specific virulence factors. The high concordance between mouse and zebrafish results reinforces the conservation of infection mechanisms in both hosts; this suggests that the three genes still untested for virulence in mice (*APM1*, *PEP8* and *NMD5*) are most likely of importance in murine (and human) disease.

The zebrafish has the unique advantage of allowing intravital observation of the early innate immune response, which enabled us to group the hypovirulent mutants into four classes. As expected, our screen revealed that increases in phagocytosis efficiency were associated with lower virulence for most of the mutants (Classes I, III and IV). For some mutant infections, phagocyte numbers were unchanged or decreased, while fungal phagocytosis levels of the mutants matched or exceeded those of wildtype cells (Classes I, II and III). In these infections, a more robust rapid phagocytosis response may limit later phagocyte recruitment at 4-6 hpi and ultimately result in lower inflammatory gene expression at later timepoints. Consistent with this idea, highly effective phagocytosis correlates with reduced epithelial NF-kB activation during mucosal *Candida* infection, which is also in line with the lower cytokine production found here at 24 hpi (58).

The cell wall and fungal morphology regulate phagocytosis by macrophages and neutrophils, with β-glucan masking and filamentous shape leading to impaired phagocytosis (9, 42). *BRG1* and *PEP8*, both in the Morphogenesis class, have not previously been identified as regulating immune responses, although both are linked to *Candida* virulence. The mechanisms underlying the roles of these genes in regulating early immune responses are unknown, although their putative functions in vesicle transport and biofilm formation both have connections to filamentous growth and surface adhesion proteins that may limit phagocytosis (40, 42, 44, 47, 59). Both mutants produced fewer elongated cells *in vivo*. Since larger filamentous cells are engulfed less efficiently, these minor deficiencies in elongation could lead to earlier fungal phagocytosis (42). These and other mutants may also have alterations in their cell walls that eliminate structural mechanisms for phagocytic evasion, although the only strains with known cell wall defects are *cek1*Δ/Δ, which has more β-glucan exposure and recruits more phagocytes to the infection site, and *rim101*Δ/Δ, which regulates cell wall genes (9, 60, 61). This conservation of phenotypes again suggests that the zebrafish is a good model for examining the effect of altered cell wall on early phagocytosis and immune recruitment—even if zebrafish do not have a direct sequence homolog of the key pattern recognition receptor for exposed β-glucan, Dectin-1 (62).

The most pronounced increase in early phagocytic efficiency occurred in infections of fungi lacking *NMD5*, a mutant in the Infectivity class which is predicted to regulate nuclear protein import and ionic stresses, based on work in baker’s yeast (47, 48). The *C. albicans nmd5*Δ/Δ mutant has defects in white-opaque switching and its expression is altered in phagocyte interaction, biofilm growth and osmotic stress (63–69). Its differential expression upon neutrophil and macrophage challenges is consistent with its role in limiting phagocytosis (65, 66, 68, 69). In contrast to its function in baker’s yeast, the *C. albicans nmd5*Δ/Δ mutant is not hypersensitive to stress conditions, suggesting a significant divergence in gene function between the species—as has been observed previously (70, 71). Instead, the function of *CaNMD5* is clearly related to immune evasion—the mutant loses its virulence disadvantage when the innate immune response is compromised by any of three methods. Given the likely role of Nmd5p in nuclear import of transcription factors, it will be interesting to identify differential transcription patterns in this mutant that may account for the loss in immune evasion.

Overall, our findings reinforce the relevance of studying *Candida*-innate immune events in zebrafish by intravital imaging, identify several new hypovirulent mutants, describe early immune evasion-related phenotypes for all of the mutants, and characterize *NMD5* as a new and important virulence factor required to limit innate immune phagocytosis. These results highlight the importance of an effective early innate immune response that engulfs *C. albicans* cells rapidly to limit germination during infection. We expect that future intravital timelapse experiments in the zebrafish at high spatio-temporal resolution will further characterize how phagocytes interact with these mutants and thereby shed light on conserved mechanisms that regulate early events in candidiasis in vertebrate hosts.

## Methods

### *C. albicans* strains and growth conditions

*C. albicans* mutant strains for screening were obtained from the Noble library (31). For infection, strains were grown on yeast-peptone-dextrose (YPD) agar at 30°C (20 g/L glucose, 20 g/L peptone, 10 g/L yeast extract, 20 g/L agar, Difco, Livonia, MI). Single colonies were picked from plates and inoculated into 5mL liquid YPD and grown overnight on a wheel at 30°C. Overnight cultures were resuspended in PBS (phosphate buffered saline, 5 mM sodium chloride, 0.174 mM potassium chloride, 0.33 mM calcium chloride, 0.332 mM magnesium sulfate, 2 mM HEPES in Nanopure water, pH = 7) and stained with Calcofluor white (750 µg/ml) when necessary. Cultures were washed twice with PBS and the concentration was adjusted to 1×10^7^ CFU/ml in PBS for injection. For imaging, strains were transformed with pENO1-iRFP-NAT^r^ according to (41). Strains were screened by fluorescence microscopy and flow-cytometry to pick the brightest isolates, and the integration site at the *ENO1* locus was confirmed by PCR as described (41). Full deletion of *RBT1* from SN250 was achieved using the SAT-flipper method as described previously (72) using LiAC transformation. The deletion cassette was generated by integrating 514 bp up and 485 bp downstream of *RBT1* into a pSFS2 derivative (72) and was excised by restriction digest with KpnI and SacI.

### Complementation of mutant strains

Complementation constructs were ordered from Genscript (Piscataway, NJ) in the pUC57 backbone and contain the ORF with 200 bp upstream and 50 bp downstream, followed by *C. dubliniensis ARG4* (Fig. S6A). Restriction sites were eliminated from the ORF during gene synthesis. A restriction site was designed within the 200 bp upstream region, an NdeI cutsite at the start of the ORF, a BamHI restriction site in *ARG4* upstream region, and a BglII site in the downstream *ARG4* region. An *NMD5* complementation construct was ordered from Twist Bioscience (South San Francisco, CA) without *ARG4.* This construct included an upstream XbaI restriction site, a 200 bp *NMD5* upstream region containing an XhoI restriction site, the *NMD5* ORF, the mNeon ORF (73) flanked by NcoI restriction sites and a PacI restriction site, then 50 bp of the *NMD5* downstream region, and a BamHI site in an *ARG4* upstream region. This region was then cloned into the Genscript pUC57 backbone by cutting with the with XbaI and BamHI to remove the *PEP8* region and replace it with the *NMD5* region to get an *NMD5* construct containing *ARG4* (Fig. S6B). For complementation, constructs were cut with the appropriate restriction enzymes, and a LiAC transformation was performed using rescue of the *ARG4* autotrophy as a selection marker. PCR was performed to ensure correct integration. *NMD5* complementation colonies were screened by flow cytometry for mNeon-positive cells. Sequences of the complementation constructs are provided in Table S5. To check for functional complementation of mutants that have known morphogenesis defect (Fig. S1) we assessed growth on Spider media (for *BRG1*, *CEK1*, and *PEP8* strains) or in M199 at pH 4 and pH 8 (for *RIM101* strains). Briefly, to test growth on Spider media, we grew SN250, mutant, and complemented strains overnight at 30°C in 5ml YPD. Overnight cultures were diluted in PBS and 100 μl of 1×10^2^ cells/ml was spread onto Spider plates. Plates were incubated at 30°C and imaged after 7 and 14 days of growth. For *RIM101* strains SN250, *rim101*/Δ/Δ, and complemented strains were grown overnight at 30°C in 5 ml YPD. 50 μl of overnight culture was inoculated into M199 pH 4 and pH 8 and grown at 37°C for 4 hours. Strains were then imaged on a Zeiss Axio Observer Z1 microscope (Carl Zeiss Microimaging, Thornwood, NJ) to assess filamentous growth.

### Growth of *nmd5*Δ/Δ on different media to assess salt tolerance

Overnight cultures were grown at 30°C in 5 ml YPD. 3×10^7^ cells from the overnight culture was inoculated into 5 ml fresh YPD and incubated on roller drum for 4 hours. After 4 hours, 10-fold serial dilutions were performed out to 10^-5^ in PBS, and 3 μl of the 10^0^ to 10^-5^ dilutions was spotted onto plates. Plates include YPD, M199 pH 8, M199 pH 4, YPD + 400 mM NaCl, YPD + 1.5 mM H_2_O_2_, YPD + 400 mM CaCl_2_, YPD + 150 mM LiCl, and YPD + 6 mM MnCl_2_. Plates were incubated at 30°C for 48 hours and imaged after 24- and 48-hours incubation. Strains were spotted in duplicate on two plates and 3 replicates performed.

### Zebrafish Care and Maintenance

Adult zebrafish were held in the University of Maine Zebrafish facility at 28°C in a recirculating system (Aquatic Habitats, Apopka Fl) under a 14 hr/10 hr light/dark cycle and fed Hikari micropellets (catalogue number HK40; Pentair Aquatic Ecosystems).

### Ethics Statement

All zebrafish studies were carried out in accordance with the recommendations in the Guide for the Care and Use of Laboratory Animals of the National Research Council (77). All animals were treated in a humane manner and euthanized with Tricaine overdose according to guidelines of the University of Maine Institutional Animal Care and Use Committee (IACUC) as detailed in protocols A2015-11-03, A2018-10-01 and A2021-09-01.

### Zebrafish Infections

Zebrafish were raised at 33°C for the first 24 hours, in E3 plus 0.3 mg/L methylene blue for the first 6 hours then E3 plus PTU (0.02 mg/ml, Sigma-Aldrich, St. Louis, Missouri) thereafter. At 24 hpf, embryos were dechorionated. Injection solutions were made up at 1×10^7^ cells/ml in PBS and stained with Calcofluor white (750 μg/mL) as necessary to visualize non-fluorescent or far-red candida by eye. Embryos were anesthetized in tricaine (160 μg/ml; Tricaine; Western Chemicals, Inc., Ferndale, WA) at the prim-25 stage for both hindbrain and yolk infection (54). Embryos that were injured during the injection process were removed. After infection fish were placed at 30°C for the remainder of the experiment and monitored for survival out to 72 hpi. Fish were screened after injection on a Zeiss Axio Observer Z1 microscope (Carl Zeiss Microimaging, Thornwood, NJ) to ensure that they received between 10-25 *C. albicans* cells.

For <u>large scale virulence screening</u>, 5 *C. albicans* mutants were tested along with SN250 WT control and PBS mock infected fish in one experiment with approximately 50 fish per strain. Due to the large number of injected fish, fish were not screened after injection and *C. albicans* was not stained with Calcofluor white. As another check, if survival of SN250 infected fish fell outside of 5.3-72.18% survival (by 72 hpi) the experiment was eliminated from consideration, and all mutant strains were retested.

### Dexamethasone treatment

For dexamethasone experiments, dexamethasone (Millipore Sigma, Calbiochem, 10 mg/ml stock) or DMSO (vehicle control) was added to the E3+PTU one hour before infection and maintained throughout the experiment. A final concentration of 50 μg/ml dexamethasone, 0.5% DMSO was used. Injections in the hindbrain ventricle were otherwise performed as described above.

### Morpholino injection

Embryos were injected between the 1- and 4- cell stage with standard (1 ng/nl, CCTCTTACCTCAGTTACAATTTATA) or p47*^phox^* (2.5 ng/nl, CGGCGAGATGAAGTGTGTGAGCGAG) morpholinos. Morpholino injection solutions were prepared in 0.3x Danieau buffer (17.4 mM NaCl, 0.21 mM KCl, 0.12 mM MgSO_4_•7H_2_O, 0.18 mM Ca(NO_3_)_2_, 1.5 mM Hepes pH 7.2) with 0.16% fluorescent dextran and 0.001% phenol red. Morpholino injected fish were kept in plain E3 until 12 hpf, when methylene blue and PTU (0.02 mg/ml) was added. At time of infection, fish were switched to E3+PTU and maintained in E3+PTU throughout infection. Fish were screened for dextran incorporation and discarded from the experiment if they did not show fluorescence throughout the fish. Hindbrain infections in morphant fish were performed at the prim-25 stage as described above.

### Quantitative real-time PCR

Fish were infected as described above, screened for correct inoculum (10-25 fungal cells), and euthanized at 4 hpi or 24 hpi for qPCR. Pools of 5-10 larvae were homogenized in TRIzol (Invitrogen, Carlsbad, CA) and stored at −80°C. RNA isolation was performed using the Direct-zol RNA MinipPrep kit (Zymo Research, Irvine, CA) following their protocol. cDNA was synthesized from 500 ng of RNA using iSCRIPT reverse transcription (RT) supermix for RT-qPCR (Bio-Rad, Hercules, CA). qPCR was performed using SsoAdvanced Universal SYBR Green Supermix (Bio-Rad) with 1 μl of cDNA in 10 μl reactions with primers listed in the table below. qPCR was run on a CFX96 Real time system, C1000 touch thermal cycler (Bio-Rad).

### Fluorescence Microscopy

For analysis of the phagocyte response at 4-6 hpi, embryos were placed in 0.4% low melting point agarose in E3 with 160 μg/ml tricaine in a glass bottom 24-well plate (MatTek Corporation, Ashland, MA) and the hindbrain ventricle imaged. Images were taken on an Olympus IX-81 inverted microscope with an FV-1000 laser scanning confocal system (Olympus, Waltham, MA) with a 20x (0.75 NA) objective with 5 μm increments for approximately 25-35 slices.

### Image Analysis

Images were imported into Fiji (ImageJ) and made into composite 4-channel z-stacks for quantification. Number of *mpeg1*:GFP+ or *lysC*:dsRed+ cells were counted manually for the hindbrain region throughout the z-stack. In addition, *C. albicans* cells were manually counted for whether they were intracellular (inside *mpeg1*:GFP+, *lysC*:dsRed+, or other), or extracellular to determine the percent of *C. albicans* cells that were taken up by the host. The total number of cells recruited to the infection included *mpeg1*:GFP+ cells, *lysC*:dsRed+ cells, as well as non-fluorescent cells phagocytosing *Candida*. Fish were excluded from the total cells recruited count if they did not contain both GFP+ and dsRed+ cells.

### Statistical Analysis

Statistical analysis was performed using GraphPad Prism software. To calculate the z-score for quantifying screening results, we measured the mean and standard deviation of 72 hpi percent survival for all of the SN250 (control) infections and then calculated [(Survival % Mutant) – (Mean Survival % Control)] / Standard Deviation of Survival % Control. For analysis of survival in non-screen experiments, Kaplan-Meier curves were generated from at least 3 pooled experiments with the same mutant *C. albicans* strains, with SN250 always included in the same experiments, and Mantel-Cox log rank tests were performed. We utilized Bonferroni corrections to reduce the family-wide error rate in exploratory experiments, while omitting this for any hypotheses that were firmly established *a priori* based on data prior to these experiments (83). Non-exploratory hypotheses based on data shown in Fig. 3 were the following: SN250 is more virulent than *nmd5Δ/Δ* but less virulent than *nmd5Δ/Δ*+*NMD5*, while *nmd5Δ/Δ* is less virulent than both the other strains. Furthermore, we have shown in previous work that p47 morpholino knockdown makes zebrafish more susceptible to wildtype *C. albicans*, so this is confirmatory rather than exploratory (30). Thus, in Fig. 7 the pairwise comparisons shown by arcs (e.g. SN250 vs. *nmd5Δ/Δ*) were not Bonferroni corrected for multiple comparisons because the effects of genotype alone were already tested in Fig. 3 and the effect of the p47 MO was already demonstrated (30). For analysis of differences in phagocyte recruitment and phagocytosis, a normality test was performed. If the distribution was not normal, the data was trimmed for outliers (top and bottom 10%) and this allowed for parametric testing. All mutants were compared with wildtype SN250 in each experiment. For simplicity to present all data in one graph, data was normalized to WT, SN250 values. For normalization, the average SN250 value for a set of experiments was divided by the average SN250 value for all experiments, to get an adjustment value. The value for each individual fish was then divided by this adjustment value, to get a normalized value for each fish. Normalized values were used to generate plots, which show the mean and 95% confidence interval. Effect size was determined as described by (84) using the Effect Size Calculator (https://f.hubspotusercontent30.net/hubfs/5191137/attachments/ebe/EffectSizeCalculator.xls). A size of greater than 0 and less than 0.3 was qualified as Small, greater than 0.3 and less than 0.5 as Moderate, and greater than 0.5 as Strong (84). Briefly, this is calculated as (M_1_ – M_2_)/ s_pooled_, where M_1_ -M_2_ is the difference between the means and s_pooled_ is the root mean squared of the two standard deviations. Hedges’ factor is used to correct for bias in effect size (85).

## Acknowledgements

We would like to thank Prof. Suzanne Noble for making the mutant library available through the Fungal Genetics Stock Center, and Mark Nilan for providing wonderful fish care. This work is supported by grants from the University of Maine Institute of Medicine (to BAB), Maine INBRE (Honors Pre-thesis and Thesis Fellowships through NIH P20GM103423 to EB, MM and LS), and R15AI169393 (to RTW). RTW is a Burroughs Welcome Fund Investigator in the Pathogenesis of Infectious Disease.

## Supplementary Figures

**Fig. S1. Complementation partially restores *in vitro* phenotypes of *brg1*Δ/Δ, *pep8*Δ/Δ, *cek1*Δ/Δ and *rim101*Δ/Δ mutants.** Growth of SN250 (**A-C)**, *brg1*Δ/Δ, *brg1*Δ/Δ+*BRG1* (**A),** *pep8*Δ/Δ, *pep8*Δ/Δ+*PEP8* (**B),** *cek1*Δ/Δ, and *cek1*Δ/Δ+*CEK1* (**C)** on Spider agar after 7 and 14 days at 30°C. **D)** Growth of SN250, *rim101*Δ/Δ, and *rim101*Δ/Δ+*RIM101* in M199 pH 4 or pH 8 after 4 hours at 37°C. Scale bar is 20 µm.

**Fig. S2. Complementation did not restore virulence of *cht2*Δ/Δ, *orf19.5547*Δ/Δ, or *rbt1^968-2166^*Δ/Δ. A)** Kaplan-Meier survival curve of fish injected with PBS (mock, n=23), SN250 (WT, n=41), *cht2*Δ/Δ (n=31), or *cht2*Δ/Δ+*CHT2* (n=44). Data pooled from 2 experiments. **B)** Kaplan-Meier survival curve of fish injected with PBS (mock, n=10), SN250 (WT, n=21), *orf19.5547*Δ/Δ (n=16), or *orf19.5547*Δ/Δ+*ORF19.5547* (n=19). Data from 1 experiment. **C)** Kaplan-Meier survival curve of fish injected with PBS (mock, n=58), SN250 (WT, n=84), *rbt1^968-2166^*Δ/Δ (n=90), or *rbt1^968-2166^*Δ/Δ+*RBT1* (n=105). Data pooled from 5 experiments. **D)** Kaplan-Meier survival curve of fish injected with PBS (mock n=35), SN250 (WT, n=60), *rbt1^968-2166^*Δ/Δ (n=52), or *rbt1*Δ/Δ (n=60). Data pooled from 3 experiments.

**Fig. S3. *brg1*Δ/Δ and *pep8*Δ/Δ show fewer elongated cells in the zebrafish hindbrain at 4-6 hours post infection. A)** Images of SN250 and *brg1*Δ/Δ infected fish showing yeast (arrow heads) and elongated cells (arrows) in the zebrafish hindbrain at 4-6 hours post infection. Scalebars are 50 µm. **B)** Plot showing the percent of elongated cells for each mutant with the control SN250 for the same experiments. The number of yeast and elongated cells was counted manually in at least 3 independent experiments for each mutant, with approximately 25 fish per strain imaged. There are separate SN250 columns for each set of experiments, as the mutant was compared to wildtype in the same experiments. Shading indicates the Groups I-IV, based on similar fungal-immune interaction phenotypes (Table 1). Means and 95% confidence intervals are plotted. Hedges bias-corrected effect sizes and significance was determined for each mutant. * indicates p<0.05, *** indicates p<0.001, # indicates a moderate effect, ## indicates a large effect.

**Fig. S4. Expression of inflammatory genes early during *C. albicans* infection.** Zebrafish larvae at the prim25 stage were infected with 10-25 *C. albicans* cells of the wildtype SN250 strain. At 4-6 hours post-infection, they were euthanized, RNA was purified, and qPCR was conducted to determine the change in gene expression relative to the mock-infected controls at the same time point. There was no significant induction of any of these inflammatory genes at this timepoint, although there was a slight reduction in *ccl2* expression. Shown are averages and 95% confidence intervals for seven biologically independent experiments. Significant changes were determined by comparing the 95% confidence intervals; none were significantly up-regulated.

**Figure S5 *<u>nmd5</u>*<u>Δ/Δ is not more susceptible to cell stressors.</u>** Growth of SN250, *nmd5*Δ/Δ, and *nmd5*Δ/Δ+*NMD5* on YPD, M199 pH 8, M199 pH 4, YPD + 400 mM NaCl, YPD + 1.5 mM H_2_O_2_, YPD + 400 mM CaCl_2_, YPD + 150 mM LiCl, and 6 mM MnCl_2_. 3×10^7^ cells from the overnight culture was inoculated into 5ml fresh YPD and incubated on roller drum for 4 hours. After 4 hours 10-fold serial dilutions 3μl of the dilutions was spotted onto plates. Plates were incubated at 30°C for 48 hours and imaged after 24- and 48-hours incubation.

**Fig. S6. Complementation Constructs A)** Plasmid showing the design of the construct for complementation of mutant strains. All plasmids contained a BglII cut site downstream of *ARG4*, a BamHI cutsite upstream of *ARG4*, an NdeI cutsite at the ORF start site, and another restriction cut site in the complementary upstream region of the gene of interest. The upstream restriction site and the BglII restriction site were used to excise the fragment for complementation. **B)** Plasmid showing the design of the construct for complementation of *nmd5***Δ/Δ**. The *NMD5* complementation construct contains mNeon to enable screening of transformants for fluorescence to assess functional complementation. The XhoI and BglII restriction sites were used to excise the fragment for complementation. Sequences of ORFs with upstream and downstream regions used in complementation constructs is provided in Table S5.

**Table S1. Full list of *C. albicans* strains used in this study.** All mutants tested in this study and which stages of testing they passed.

**Table S2. Phagocytosis efficiency for Calcofluor White-labeled mutant *C. albicans* infections.** Summary statistics of the immune response to infection for all mutant *C. albicans* infections imaged, as shown in Fig. 1E.

**Table S3. Immune response to mutant *C. albicans* infections.** Summary statistics of the immune response to infection for all mutant *C. albicans* infections imaged in double transgenic fish, as shown in Fig. 4 and Fig. 5.

**Table S4. Comparison of mutant virulence in zebrafish hindbrain infection versus mouse tail vein infection.** Breakdown of virulence phenotypes for all mutants included in this screen.

**Table S5. Complementation Construct Sequences.** Sequences used for complementation for each mutant, as well as *C. dubliniensis ARG4* used in each of the complementation constructs. Complementation constructs were constructed by inserting these sequences into the pUC57 vector.

