## Supplementary figures and images for "Forward genetic screen in zebrafish identifies new fungal regulators that limit host-protective *Candida*-innate immune interaction"

### Supplemental Fig. 1

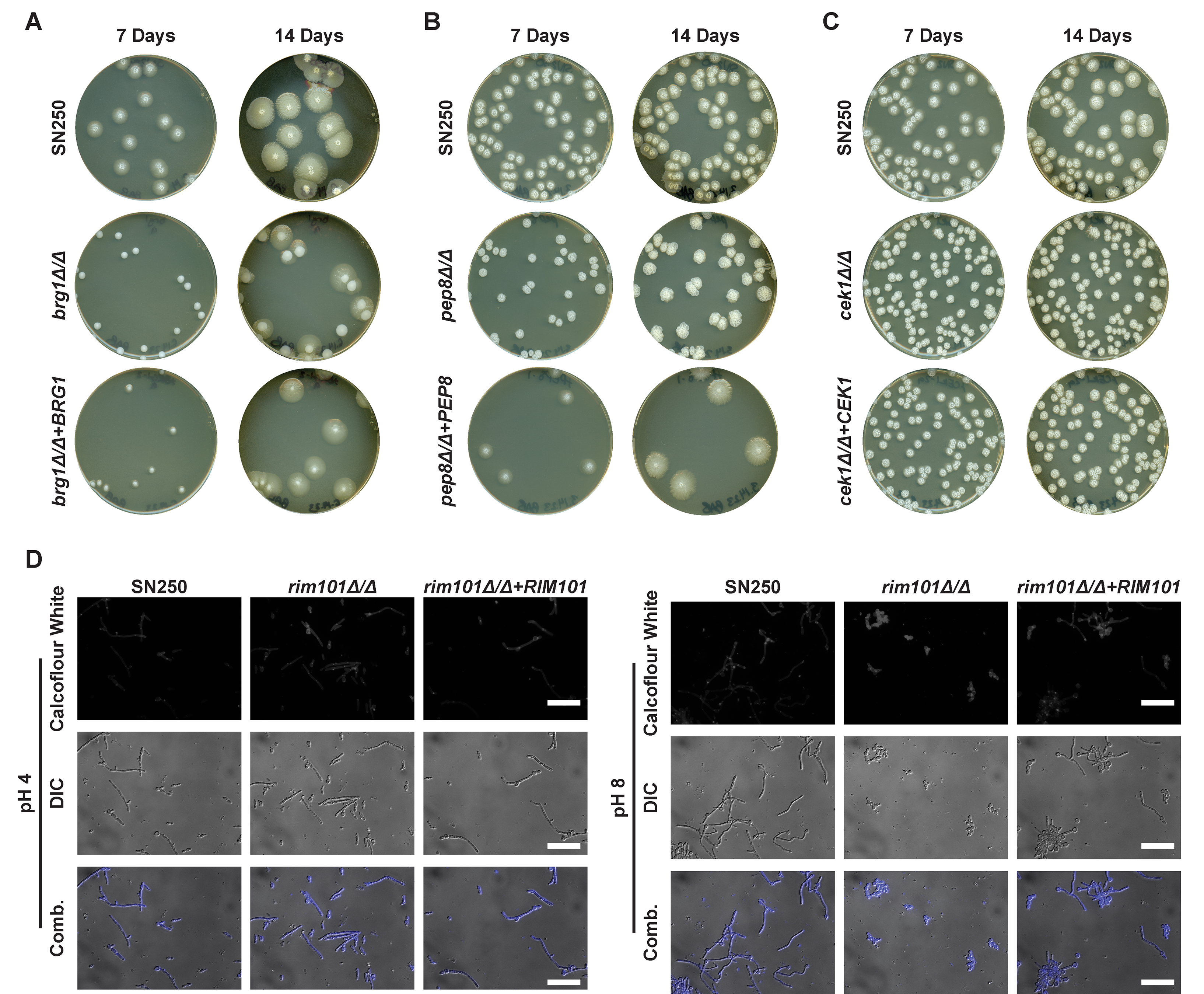

### Supplemental Fig. 2

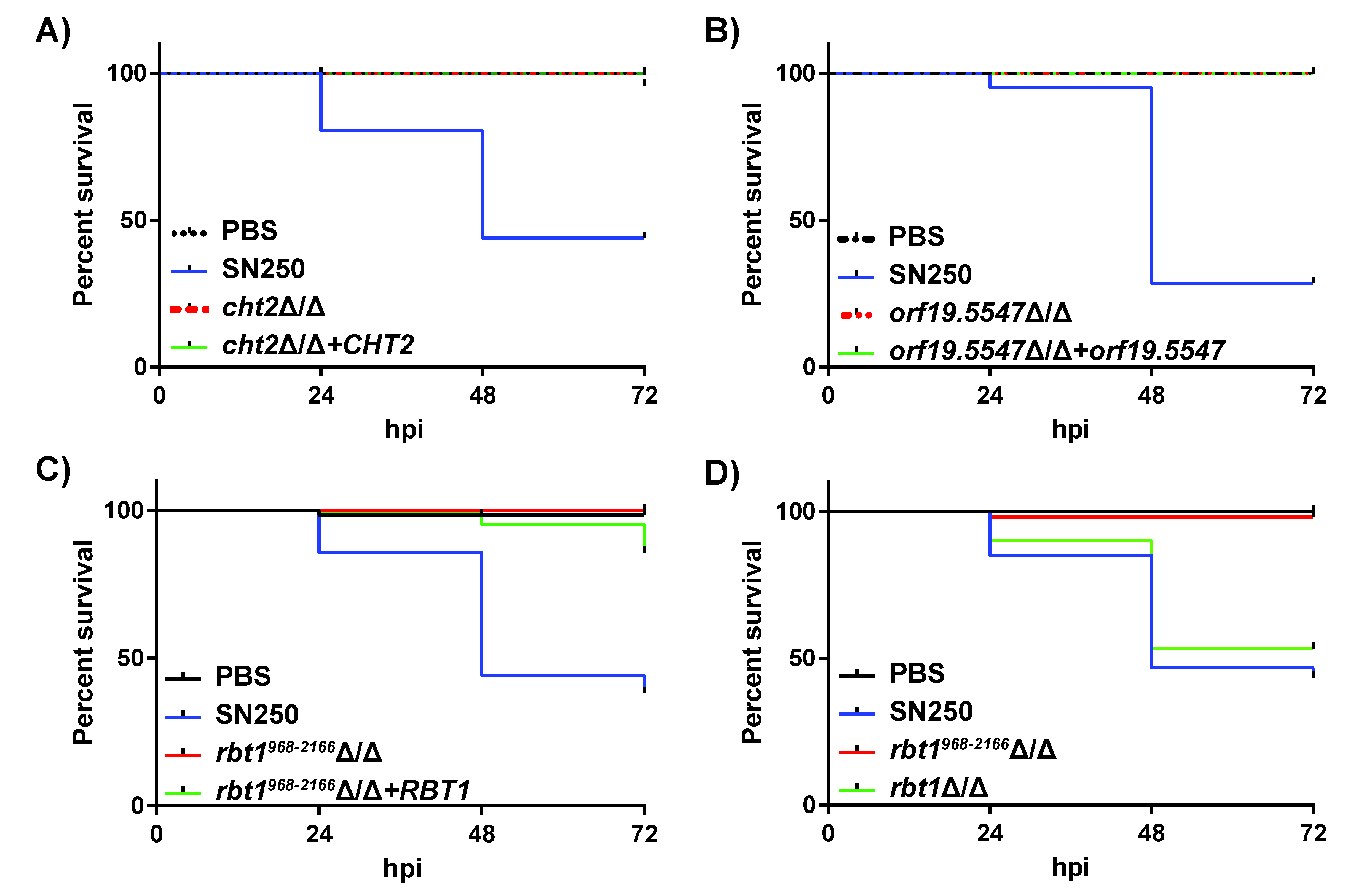

### Supplemental Fig. 3

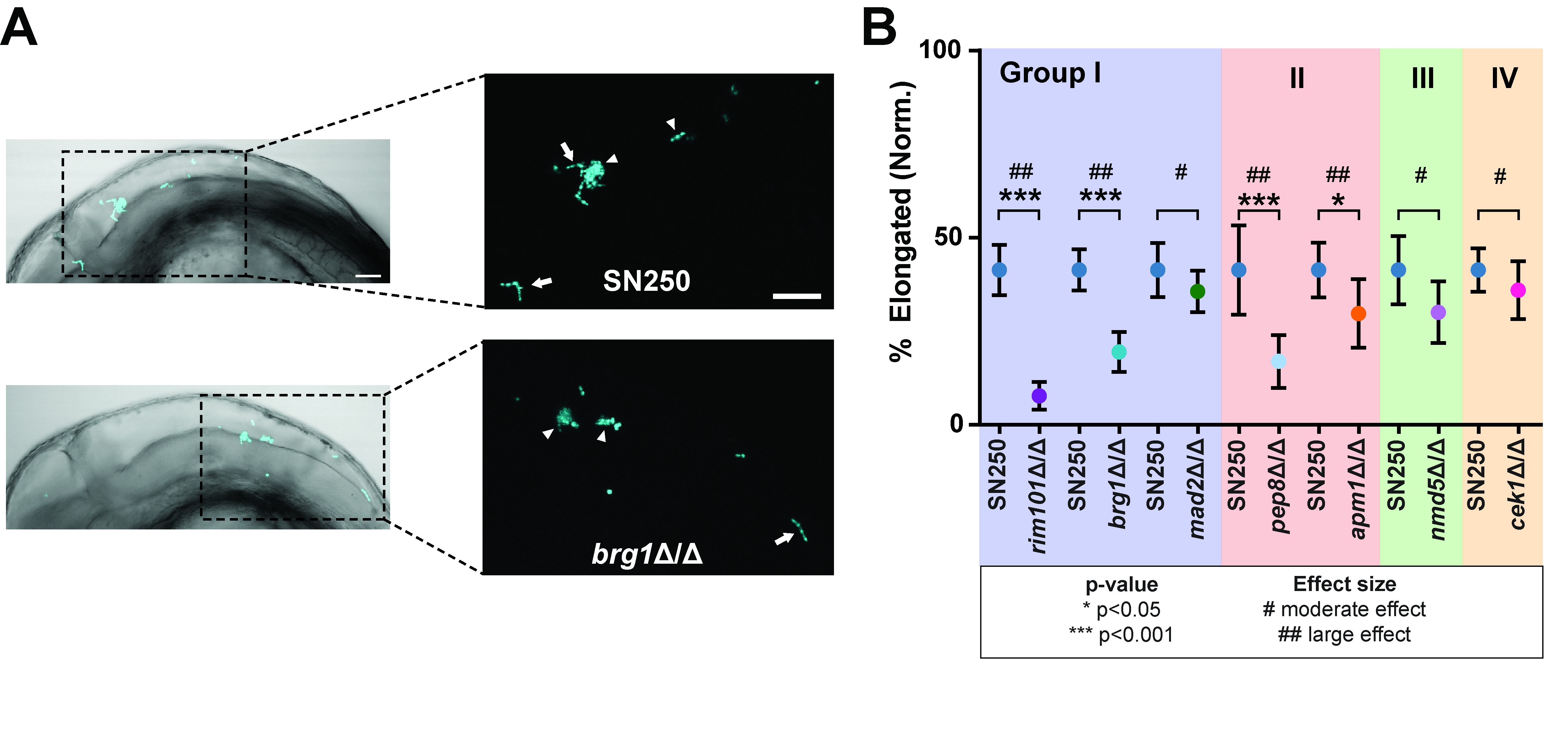

### Supplemental Fig. 4

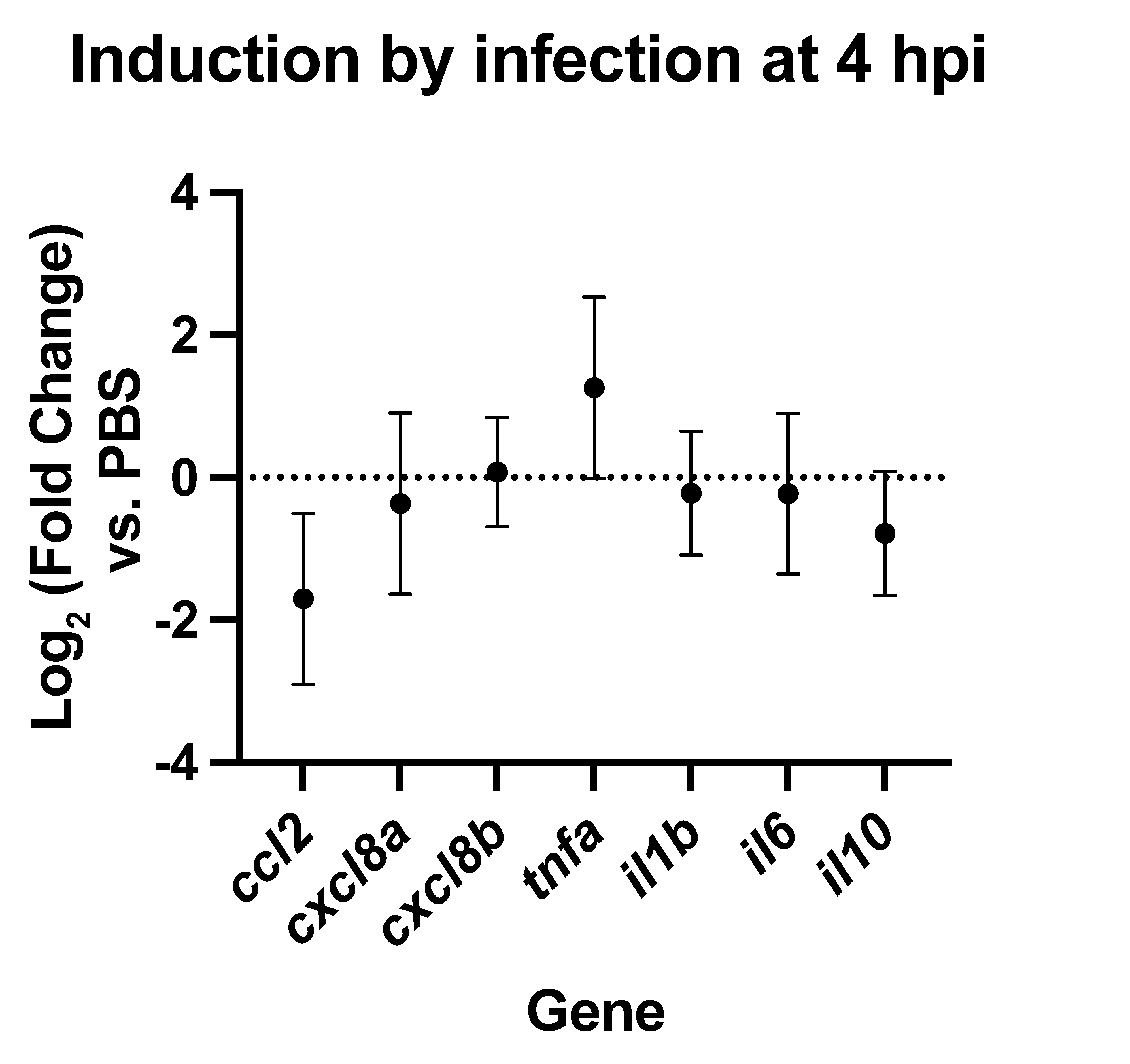

### Supplemental Fig. 5

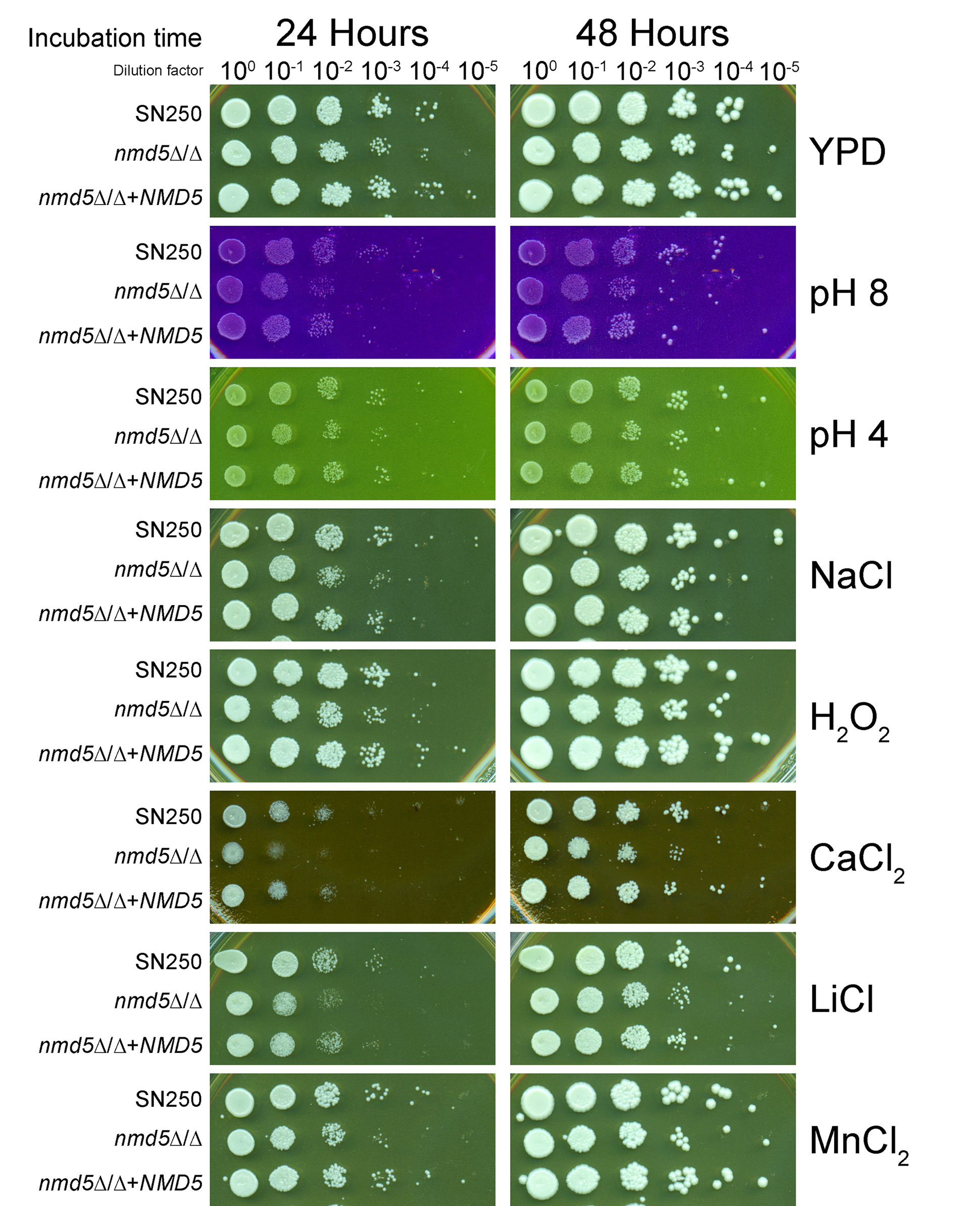

### Supplemental Fig. 6

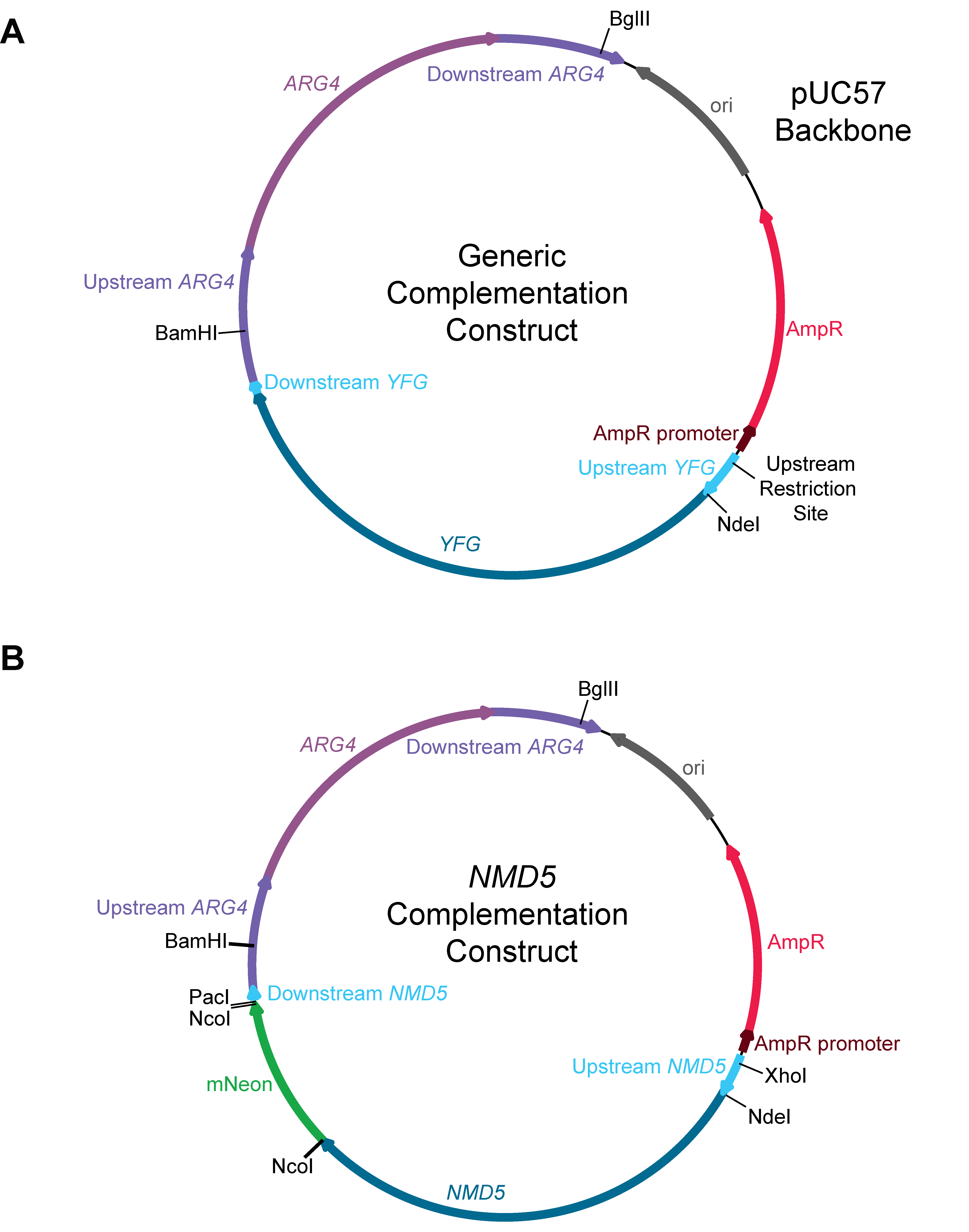
